# The impact of the Turkish (TK) population variome on the genomic architecture of rare disease traits

**DOI:** 10.1101/2020.04.27.064824

**Authors:** Zeynep Coban-Akdemir, Xiaofei Song, Francisco C. Ceballos, Davut Pehlivan, Ender Karaca, Yavuz Bayram, Tadahiro Mitani, Tomasz Gambin, Tugce Bozkurt Yozgatli, Shalini N. Jhangiani, Donna M. Muzny, Richard A. Lewis, Baylor Hopkins Center for Mendelian Genomics, Pengfei Liu, Eric Boerwinkle, Ada Hamosh, Richard A. Gibbs, V. Reid Sutton, Nara Sobreira, Claudia M. B. Carvalho, Chad A. Shaw, Jennifer E. Posey, David Valle, James R. Lupski

## Abstract

**Purpose:** The variome of the Turkish (TK) population, a population with a considerable history of admixture and consanguinity, has not been deeply investigated deeply for its potential impact on the genomic architecture of disease traits.

**Methods:** We generated and analyzed a database of variants derived from exome sequencing (ES) data of 773 TK unrelated, clinically affected individuals with various suspected Mendelian disease traits, and 643 unaffected relatives.

**Results:** Using uniform manifold approximation and projection (UMAP), we showed that the TK genomes are more similar to those of Europeans and consist of two main subpopulations: clusters 1 and 2 (N=235 and 1,181) that differ in admixture proportion and variome (https://turkishvariomedb.shinyapps.io/tvdb/). Furthermore, the higher inbreeding coefficient (*F*) values observed in the TK affected compared to unaffected individuals correlated with a larger median span of long-sized (>2.64 Mb) runs of homozygosity (ROH) regions (*p*-value=2.09e-18). We show that long-sized ROHs are more likely to be formed on recently configured haplotypes enriched for rare homozygous deleterious variants in the TK-affected compared to TK-unaffected individuals (*p*-value= 3.35e-11). Analysis of genotype-phenotype correlations reveals that genes with rare homozygous deleterious variants in long-sized ROHs provide the most comprehensive set of molecular diagnoses for the observed disease traits with a systematic quantitative analysis of HPO (Human Phenotype Ontology) terms.

**Conclusion:** Our findings support the notion that novel rare variants on newly configured haplotypes arising within the recent past generations of a family or clan contribute significantly to recessive disease traits in the TK population.

## INTRODUCTION

Population genetics has been applied for many decades as a means to uncover loci that contribute to human disease by the genome-wide analyses of single nucleotide polymorphisms (SNPs). These SNPs are common variants, frequently shared among geographically distinct populations, and enable a fine-scale mapping of recombination to yield potential disease contributing loci. This GWAS approach has been applied largely to the study of common, complex conditions for which many have hypothesized that a combination of common variants each with small effect size can influence genetic susceptibility to disease expression^1,2^.

More recent studies, however, have uncovered a rare variant contribution to some apparently complex conditions (*i.e*., chronic kidney disease, scoliosis, arthrogryposis, developmental delay, and intellectual disability), suggesting that a substantial proportion of seemingly common, complex conditions may represent in fact a combination of individually rare, Mendelian disease traits^3–7^.

The role of ultra-rare variants in recessively inherited conditions has been less studied, although novel variants contributing to recessive disease have been reported in non-European origin population-specific cohorts^8–10^. To investigate comprehensively the role of new mutations in rare recessive disease traits, we created a population-specific rare disease cohort and rare variant database against which to measure population-based frequencies in the context of population genetic substructure.

The transmission of traits, genes, and variant alleles from one generation to the next may result in identity-by-descent (IBD) at a locus in a population characterized by consanguinity or a founder effect due to a historical population bottleneck or geographic isolation. Experimentally, evidence for IBD in an individual genome is suggested by the presence of runs of homozygosity (ROH) not accompanied by copy-number variation; i.e., copy number neutral gene dosage at a locus. Analysis of ROH regions in an individual genome can be used to prioritize potential pathogenic variations at a gene locus and may unveil possible genetic susceptibility to underlying disease^11–13^.

The inter-individual variation in ROH number and total length has also been shown to contribute to the genetic architecture of complex traits and rare diseases^14–17^. Therefore, analysis of ROH regions can be an adjuvant analytical tool to address the genetic architecture of disease. For instance, homozygosity mapping has been a central and robust genetic approach for the identification of biallelic variants causing autosomal recessive (AR) disease traits. For the identified “disease gene”, the locus resides in ROH blocks shared among affected individuals^18–23^. This approach was used successfully to map disease genes in consanguineous families with heterogeneous neurological disorders^24^, multisystem disorders^25,26^, and inborn errors of metabolism^8–10^. Furthermore, analysis of rare homozygous deleterious variants in ROH regions can elucidate the genetic heterogeneity and the molecular mechanisms underlying Mendelian traits due to either the contribution of multi-locus variation or oligogenic/polygenic recessive effects ^3,7,27–33^, and even provide a molecular genetic explanation for trait penetrance, variability of expression of disease^34^, and evidence for potential modifying gene loci^35–37^.

Both the frequency and the degree of relatedness for consanguineous marriages within populations vary in different geographic regions and countries around the globe; the highest documented frequency is recorded in Pakistan, reported as 60-76%^38^ of unions. Although consanguineous families facilitate genetic locus mapping and disease gene discovery, many populations with a relatively high coefficient of consanguinity, such as the Turkish (TK) population^39,40^, have not been investigated deeply with genomic studies, including a paucity of clinical genomics studies. The population genetics of Turkey presents peculiar features compared to other Middle East countries. Besides having a high reported rate of consanguinity, 20.1%^41^, Turkey also is described often as both a geographic and a social ‘bridge’ between Asia and Europe, an important hub of both ancient and contemporary population migration. Population substructure studies in the TK population potentially can provide insights about the effects of high admixture and a relatively increased rate of consanguinity to impact the genomic architecture of disease.

To investigate the influences of both admixture and consanguinity on population genetic variation and the genetic architecture of disease, we studied a sizeable TK cohort (1,416 personal genomes) with a considerable amount of consanguinity and admixture. We performed ES and family-based genomic analysis on 1,416 TK individuals consisting of 773 unrelated clinically affected individuals with a wide variety of suspected Mendelian disease traits, and 643 unaffected relatives. We performed an unbiased exome variant analysis that would enable a rare variant family-based genomics approach to elucidate the molecular etiology and define gene loci potentially contributing to their clinically observed disease traits. We further carried out systematic genomic and phenotypic analysis of this population, by genomic variant detection and with a structured ontology of human phenotype ontology (HPO) terms, that might reveal key features of the TK population variome and substructure that affect the genetic architecture of disease in this TK cohort.

## MATERIAL AND METHODS

### Experimental model and participant details

We recruited 1,416 unrelated TK individuals (669 females and 747 males) in the Baylor Hopkins Center for Mendelian Genomics (BHCMG) cohort (data freeze: December 2011- October 2020) after all relevant subjects or legally authorized representatives provided written informed consent for the use of their DNA and personal genomes for the identification of potential disease-contributing variants and for broad data sharing. Peripheral blood was collected from affected individuals, parents, and unaffected relatives if available. Genomic DNA was extracted from blood leukocytes according to standard procedures. All genomic studies were performed on DNA samples isolated from blood. This study was approved by the Institutional Review Board at Baylor College of Medicine (IRB protocol # H-29697).

### Materials availability

This study did not generate unique biological or chemical reagents.

### Exome Sequencing (ES) and annotation

ES was performed at the Human Genome Sequencing Center (HGSC) at Baylor College of Medicine through the Baylor-Hopkins Center for Mendelian Genomics (BHCMG) initiative. With 1ug of DNA, an Illumina paired-end pre-capture library was constructed according to the manufacturer’s protocol (Illumina Multiplexing_SamplePrep_Guide_1005361_D) with modifications described in the BCM-HGSC Illumina Barcoded Paired-End Capture Library Preparation protocol. Pre-capture libraries were captured into 4-plex library pools and hybridized in solution to the HGSC-designed Core capture reagent (52Mb, NimbleGen) or 6-plex library pools with the custom VCRome 2.1 capture reagent (42Mb, NimbleGen), according to the manufacturer’s protocol (NimbleGen SeqCap EZ Exome Library SR User’s Guide) with minor revisions. The sequencing was performed in paired-end mode with the Illumina HiSeq 2000 platform or Illumina NovaSeq 6000 platforms. Data were aligned to GRCh37/hg19 with BWA-aln (for data generated on HiSeq 2000) or BWA-mem (for data generated on NovaSeq). Sequence analysis was performed with the HGSC Mercury analysis pipeline (https://www.hgsc.bcm.edu/software/mercury)^42,43^, which moves data through various analysis tools from the initial sequence generation to annotated variant calls (SNPs and intra-read insertion/deletions; i.e. indels). Variants were called with ATLAS2 or xATLAS^44^ and the Sequence Alignment/Map (SAMtools) suites, and annotated with an in-house-developed Cassandra^45^ annotation pipeline that uses Annotation of Genetic Variants (ANNOVAR)^46^ and additional tools and databases including ExAC (http://exac.broadinstitute.org), gnomAD (https://gnomad.broadinstitute.org), the GME variome (http://igm.ucsd.edu/gme/), and the ARIC database (http://drupal.cscc.unc.edu/aric/).

### Phenotypic characterization of the BHCMG cohort

We used the PhenoDB database to collect and store the information of clinical features, pedigree structures, and self-reported consanguinity levels. Computational analyses of phenotyping data were performed by HPO terms analyses described as a phenotypic similarity score with the R package ontology Similarity^29,47,48^.

### Obtaining a final set of unrelated individuals in the TK cohort

To minimize overrepresentation of individuals from the same family and to identify mutually unrelated individuals in the TK cohort, we used the PC-AiR function in the R GENESIS package, that measures pairwise kinship coefficients and ancestry divergence to identify an ancestry representative subset of mutually unrelated individuals. Removing individuals with a close relationship from the analysis resulted in a final set of 1,416 unrelated TK individuals: 773 unrelated affected and 643 unrelated unaffected participants.

### Population substructure analysis

We investigated the population structure of this TK cohort along with the African, East Asian, South Asian, and European population samples from the 1000 Genomes Project phase 3 release data including 2,504 individuals’ genotypes in total (http://ftp.1000genomes.ebi.ac.uk/vol1/ftp/release/20130502/). First, genotypes for 31 individuals who have a biological relationship with the 2504 samples were removed from the analysis. Variants falling out of the HGSC-designed Core capture design and VCRome 2.1 capture design in the 1000 Genomes Project data were filtered out from the analyses. A pruned subset of the remaining polymorphic SNVs that are in approximate linkage equilibrium of each other (N=89,379 for the TK affected participants and N=86,049 for the TK unaffected participants) was used for the Uniform manifold approximation and projection (UMAP). UMAP was performed with the umap R package. For the admixture analysis, we ran the unsupervised ADMIXTURE algorithm by k=5 clusters that outputted the minimal cross validation error.

### Estimation of inbreeding coefficient values

The coefficient of inbreeding of an individual represents the probability that two alleles at any randomly chosen locus in an individual are identical-by-descent.

The inbreeding coefficient values of the TK cohort were estimated from the ES data with plink –het function filtering the variants with MAF >= 0.05.

### Identifying and analyzing ROH segments from ES data

We detected ROH regions from unphased exome sequence (ES) data as Absence of Heterozygosity (AOH) genomic intervals with BafCalculator (https://github.com/BCM-Lupskilab/BafCalculator)^32^. To call ROH regions with BAFCalculator, we extracted all the high-quality SNVs residing in the capture region (mostly exonic regions) available from the VCF file of each single individual’s exome. For those SNVs, we extracted a B-allele frequency (i.e., variant reads/total reads ratio); then, we transformed this ratio by subtracting 0.5 and taking the absolute value for each data point. Transformed B-allele frequency data were processed with Circular Binary Segmentation (CBS) implemented in the DNAcopy R Bioconductor package^49,50^ to call the ROH regions. This algorithm merges the consecutive exon calls, and so in this way we can detect ROH regions all over the genome that are of size ranging from hundreds of Kb to a few Mb, including a number of genes. To test the false positive and false negative rate of exome data and our algorithm, BAFCalculator, we ran an independent analysis. We ran the BAFCalculator to call ROH regions from an independent dataset consisting of 929 samples with both genome sequencing (unphased GS) and high resolution phased array data available in the Human Genome Diversity panel (https://www.internationalgenome.org/data-portal/data-collection/hgdp). Then we compared ROH regions identified by the BAFCalculator with the GS data to the ROH regions detected through high resolution array in those 929 samples. The BAFCalculator algorithm was further optimized by the segmentation mean (seg.mean) parameter (an absolute measure of average homozygosity rate of a putative ROH call). Cross-referencing the ROH regions identified by the fine-tuned BAFCalculator to the array data ROH calls showed a positive predictive value of 90% and true positive rate of 72% when the seg.mean parameter=0.47 (Supplementary Figure 1). After this analysis, we also took the exome portions of the genome data in those samples and ran the BAFCalculator with only those regions. Then, we compared FROH (the total size of ROHs >= 1.5 Mb) estimates identified from those regions (FROH_WES) to the FROH estimates identified from the genome sequencing data (FROH_WGS). These analyses revealed that, when the seg.mean=0.47, this provides a nearly perfect correlation (0.98) between FROH_WES and FROH_WGS, and that ES performs similarly to GS (Supplementary Figure 2). In summary, segments with the mean signal > 0.47 and number of marks >= 10 were classified as ROH regions.

The calculated ROH intervals from BafCalculator could represent individual genomic/gene loci resulting in ROH for diploid alleles that can occur by: i) IBD, ii) UPD^51^, or iii) a large deletion CNV. To exclude the ROH blocks that could be caused by genomic overlapping of a common variant deletion CNV, we first identified deletion CNVs through XHMM^52^. We further intersected ROH segments and potential deletion CNVs with BEDTools ^53^, and then retained only ROH regions overlapping less than 50% of their size with a variant deletion CNV. We grouped ROH regions into three size categories, applying Gaussian-mixture modeling from the MClust function in mclust R package into three length classes: long-sized genomic intervals or ROH blocks (>2.64 Mb), medium-sized ROH blocks (0.671-2.64 Mb), and short ROH blocks (0.210-0.671 Mb).

To control for the variable mutation rates across different genomic regions and among different individuals, for each individual *i* and ROH region category *r*, *r*, ∈ {*total* – *ROH*, *non* – *ROH*, *long*–*sized ROH*, *medium*–s*ized ROH, short*–*sized ROH*, we computed a variant density *f_ir_*

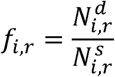

in which 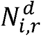 is the count of rare homozygous deleterious or likely damaging variant alleles (variants above a certain CADD score (>=15) that are located within a region *r*, while 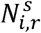 is the count of synonymous variants in a region *r*.

### Generation of an objective score for phenotypic similarity comparison

We applied two R packages, OntologyIndex and ontologySimilarity, to measure the phenotypic similarity between two sets of human phenotype ontology (HPO) terms; each set of terms was associated with a patient’s clinical features recorded in PhenoDB (https://phenodb.org/)^54^. To assess the ability of a ‘candidate disease-contributing gene’ to explain a patient’s clinical features, we applied a previously published method, in which first a MICA (most informative common ancestor) matrix was calculated for each pair of HPO terms, and then a Resnik score was calculated for two sets of HPO terms^29,47,48^.

First, we calculated the information content (IC) for HPO term *t* by

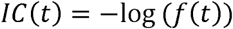

Where *f*(*t*) is the frequency of term *t* observed in all of the OMIM entries. The similarity of term *i* and term *j* is calculated with a Resnik method:

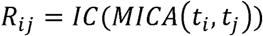

where the Resnik score, *R_ij_*, is determined by the IC of the most informative common ancestor (MICA) of term *i* and term *j*. Next, we defined the phenotypic similarity score *Sim* for two HPO sets *l*_1_ and *l*_2_ as follows:

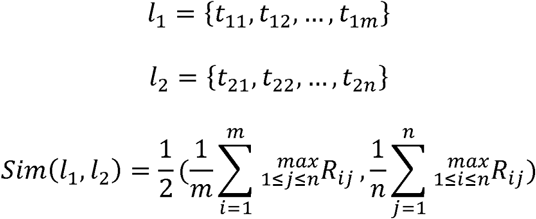

The main features of known human disease genes were summarized in OMIM (https://www.omim.org/) in the format of both plain texts and clinical synopses, and the associated HPO terms for each OMIM entry were annotated manually by the HPO-team (https://hpo.jax.org/app/download/annotation). We adapted this method to measure the phenotypic similarity between a patient’s phenotypes and a list of disease genes. For each list of the tested disease genes, we calculated a z-score performing 1,000 simulations; in each simulation we computed a similarity score between the patient’s clinical features and the associated HPO terms of a randomly selected disease gene list, which has a same number of genes as the tested disease gene list. Further, to compare the contribution of disease gene lists across different genomic regions to the explanation of a patient’s clinical phenotypic features, we computed a ratio of the similarity score calculated for a subset of disease genes, e.g., genes located in long-sized ROH regions to the similarity score calculated for all the associated disease genes in a patient’s genome.

### Statistical Analyses

We performed the statistical analyses with R version 3.3.3. We compared pairwise differences in the average values of estimated inbreeding coefficient values (F), homozygous rare deleterious variant burden (density) and total, median length, and count of ROHs in two participant groups, TK affected and TK unaffected participants by the Wilcoxon-rank sum one-tailed test. Bar plots, box plots, pie charts, and scatter plots were generated by ggplot2 data visualization R package, and stat_pvalue_manual function in the ggpubr R package added P-values and significance levels to those plots. Ddply function in plyr CRAN R package reported the summary statistics of estimated F values, homozygous rare deleterious variant burden (density) and total, median length and count of ROHs in 2 participant groups, TK affected and TK unaffected participants.

## RESULTS

### The fine-scale population substructure of the TK cohort

To study finer-scale population substructure of the TK individuals in comparison to the African, East Asian, European, and South Asian population samples from the 1000 Genomes Project, we performed the uniform manifold approximation and projection (UMAP) dimension reduction method. The first and second main UMAP components separated the samples from African, East Asian, European, South Asian, and TK populations. These studies showed that the TK genomes were distinct from the African, East Asian, and South Asian populations, but closely clustered with the variome of European samples and consist of two main subpopulations (cluster 1 (N=235) and cluster 2 (N=1,181) individuals, Figure 1A and B).

**Figure 1.**
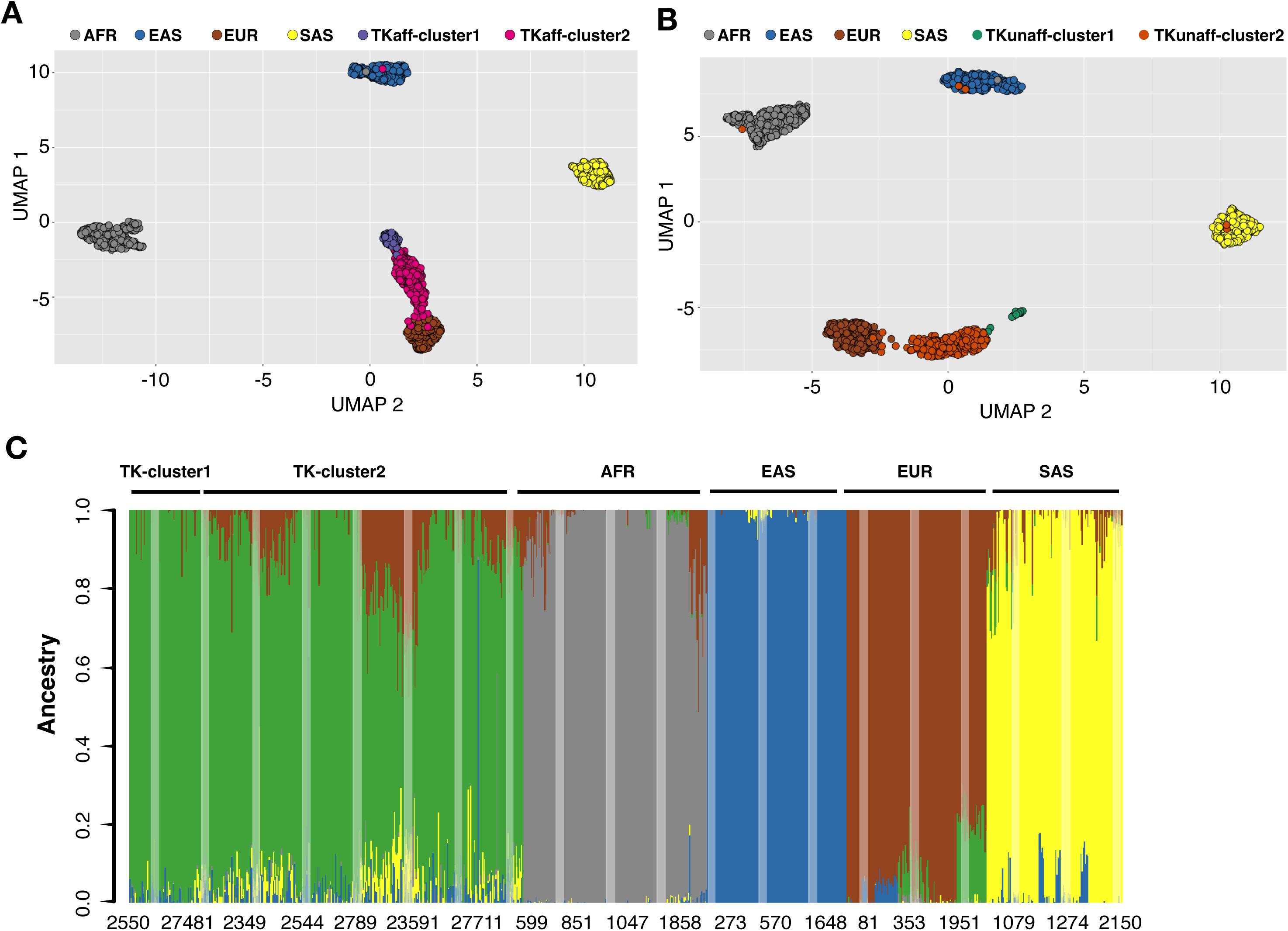
Fine-scale population substructure of TK cohort. Scatter plots display uniform manifold approximation and projection (UMAP) analysis that compares the population structure among the **A)** TK affected participants (N=773, colored in purple and dark pink) and **B)** TK unaffected participants (N=643, colored in dark green and dark orange) of the BHCMG cohort to the African (AFR) (N=661, colored in gray), East Asian (EAS) (N=504, dark blue), European (EUR) (N=503, brown) and South Asian (SAS) (N=489, yellow) population samples from the 1000 Genomes project. **C)** The bar plot demonstrates the results of the admixture analysis performed through the unsupervised ADMIXTURE algorithm (k=5 clusters) quantifying the fraction of ancestry proportions contributed by the African (gray), East Asian (dark blue), European (brown), South Asian (yellow) and Other (pink) for the TK cluster 1 and cluster 2 individuals.

To evaluate the population substructure of the TK individuals, we ran the unsupervised ADMIXTURE algorithm by k=5 clusters to minimize cross validation error. The admixture analysis results revealed that, compared to the first cluster (N=235), the second cluster (N=1,181) demonstrated a higher fraction of East Asian, European, South Asian (Wilcoxon test one-tailed P-values= 1.39e^-^4, 2.88e^-47^ and 2.36e^-25^), and a lower fraction of other (Wilcoxon test one-tailed P-values= 2.36e^-25^) ancestry, but do not differ significantly from each other in the African ancestry component (P-values = 0.172 Supplementary Figure 3).

### Comparison of the TK cohort variome to control for variant database bias

We next questioned how the population substructure was represented in the TK variome by examination of the characteristics of the distinct 3,024,990 variants present in TK individuals. We found that 2.6% (N=79,628) and 17.4% of those variants (N=527,528) are present uniquely in the TK unaffected cluster 1 or 2 individuals, and 13% (N=394,010) and 16.3% of those variants (N=493,079) are present uniquely in the TK affected cluster 1 or 2 individuals, respectively (Figure 2A). We then surveyed all variants of the TK variome in control variant databases including the genome aggregation database (gnomAD, https://gnomad.broadinstitute.org)^55^ and the Greater Middle Eastern (GME) variome^56^. Regarding the variome in the TK unaffected participants, the overall comparison of all distinct SNVs identified showed that 27% and 26% of the unique variants in cluster 1 (N=660,255 variants) and cluster 2 individuals (N=1,845,686 variants) respectively were present in the gnomAD variome. Intriguingly, only 10% and 5% of the cluster 1 and 2 variants respectively in the TK unaffected participants were present in the GME variome, underscoring the necessity of population-matched control databases to assess accurately variant minor allele frequency (Figure 2B and C and Supplementary Figure 4A and B).

**Figure 2.**
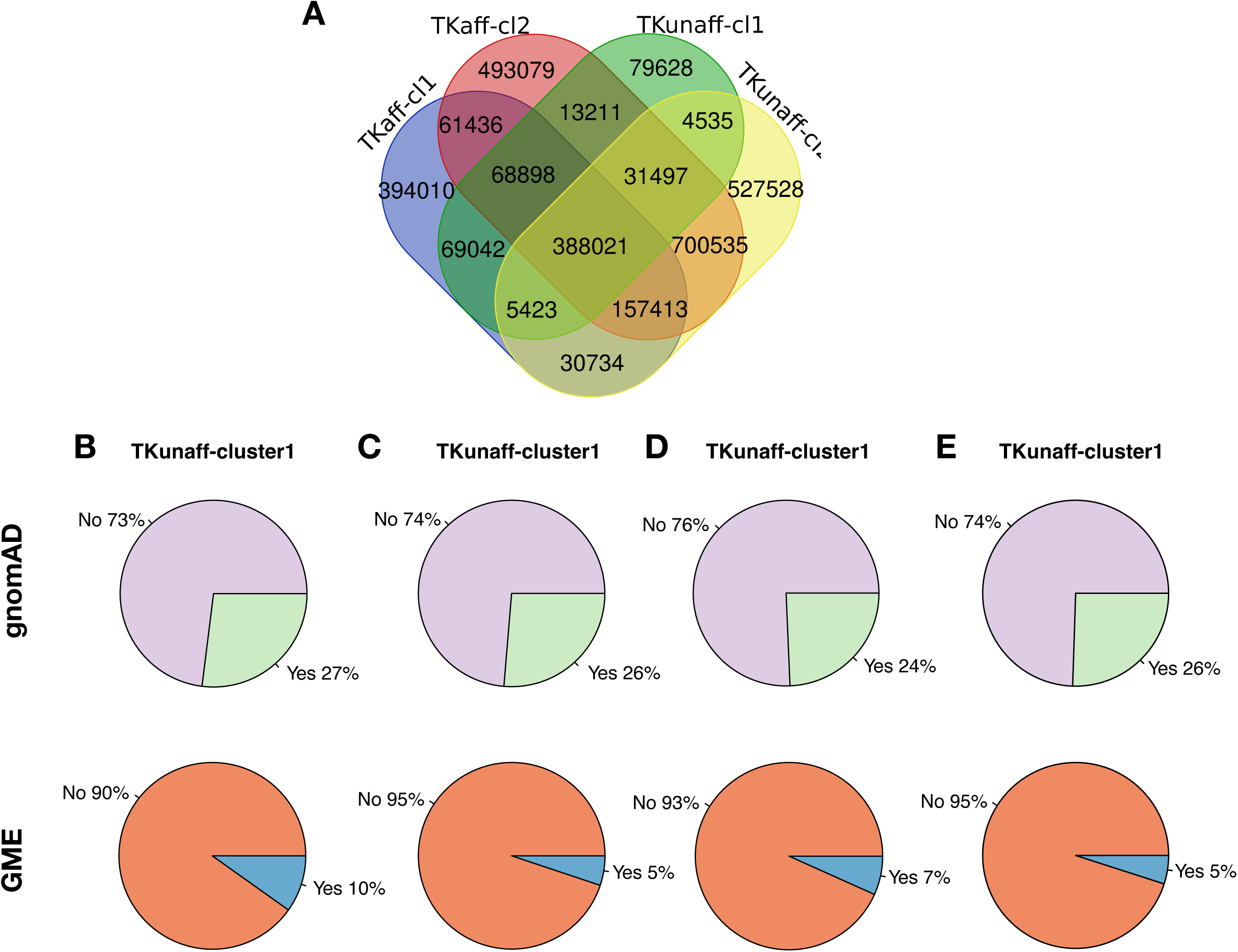
Comparison of the TK cohort variome to control databases. **A)** The Venn Diagram depicts the number of variants present in the TK affected cluster 1 (TKaff-cl1), affected cluster 2 (TKaff-cl2), unaffected cluster 1 (TKunaff-cl1) and unaffected cluster 2 (TKunaff-cl2) participants (N=3,024,990 variants in total). **B-E)** The pie charts represent all distinct variants in the TK cohorts and whether they are present (‘yes’) or absent (‘no’) in gnomAD (upper panel) and the Greater Middle Eastern Variome (lower panel) by affected status and cluster: **B)** unaffected cluster 1; **C)** unaffected cluster 2; **D)** affected cluster 1 and **E)** affected cluster 2.

We also performed similar analyses for the variome of TK affected participants (N=773). These studies demonstrated that 24% and 26% of cluster 1 (N=1,174,977 variants) and cluster 2 (N= 1,914,090) variants, respectively, were represented in the gnomAD database. Of note, only either 7% or 5% of the cluster 1 and cluster 2 variants were represented in the GME variome (Figures 2D and E and Supplementary Figure 4C and D). Aggregate TK exome variant data from clusters 1 and 2 are available publicly for analysis through a TK variome database (https://turkishvariomedb.shinyapps.io/tvdb/).

### Observation of higher estimated inbreeding coefficient values in the TK participants

To obtain a more objective experimental measure of the consanguinity level, we estimated the inbreeding coefficient (F) values from ES data of the TK affected (N=773) and unaffected participants (N=643) compared to the African (N=661), East Asian (N=504), European (n=503), and South Asian (N=489) population samples from the 1000 Genomes Project. These analyses revealed that the estimated F values in the TK unaffected participants were significantly higher with mean values of 0.017 and 0.029 in cluster 1 and cluster 2 individuals respectively, when compared to African (0.004), East Asian (-0.001), European (-0.0006), and South Asian (0.012) persons (Figure 3A). We showed further that the estimated F values of TK affected cluster 1 and cluster 2 individuals (mean=0.055 vs. 0.052) were increased significantly compared to the TK unaffected cluster 1 and cluster 2 individuals respectively (Wilcoxon test one-tailed P-values 2.3e-6 and 2.2e-16, Figure 3A and B). On the other hand, we did not observe any significant cluster-specific differences in the estimated F values of either TK affected (Wilcoxon test one-tailed P-value 0.34) or TK unaffected individuals (Wilcoxon test one-tailed P-value 0.11, Figure 3B). Therefore, we merged these two clusters for further analyses. Our analysis also showed that the measured genomic inbreeding coefficients - the fraction of the genome covered by ROHs > 1.5 Mb (FROH) - are 0.048 and 0.030 on average in the TK affected and unaffected participants respectively, and these are nearly 1-1 proportional to the average estimated F values of the TK affected [0.053] and unaffected participants [0.028]. These findings support the contention that the excess of homozygosity in the TK genomes was shaped mostly by ROHs (Figure 3C).

**Figure 3.**
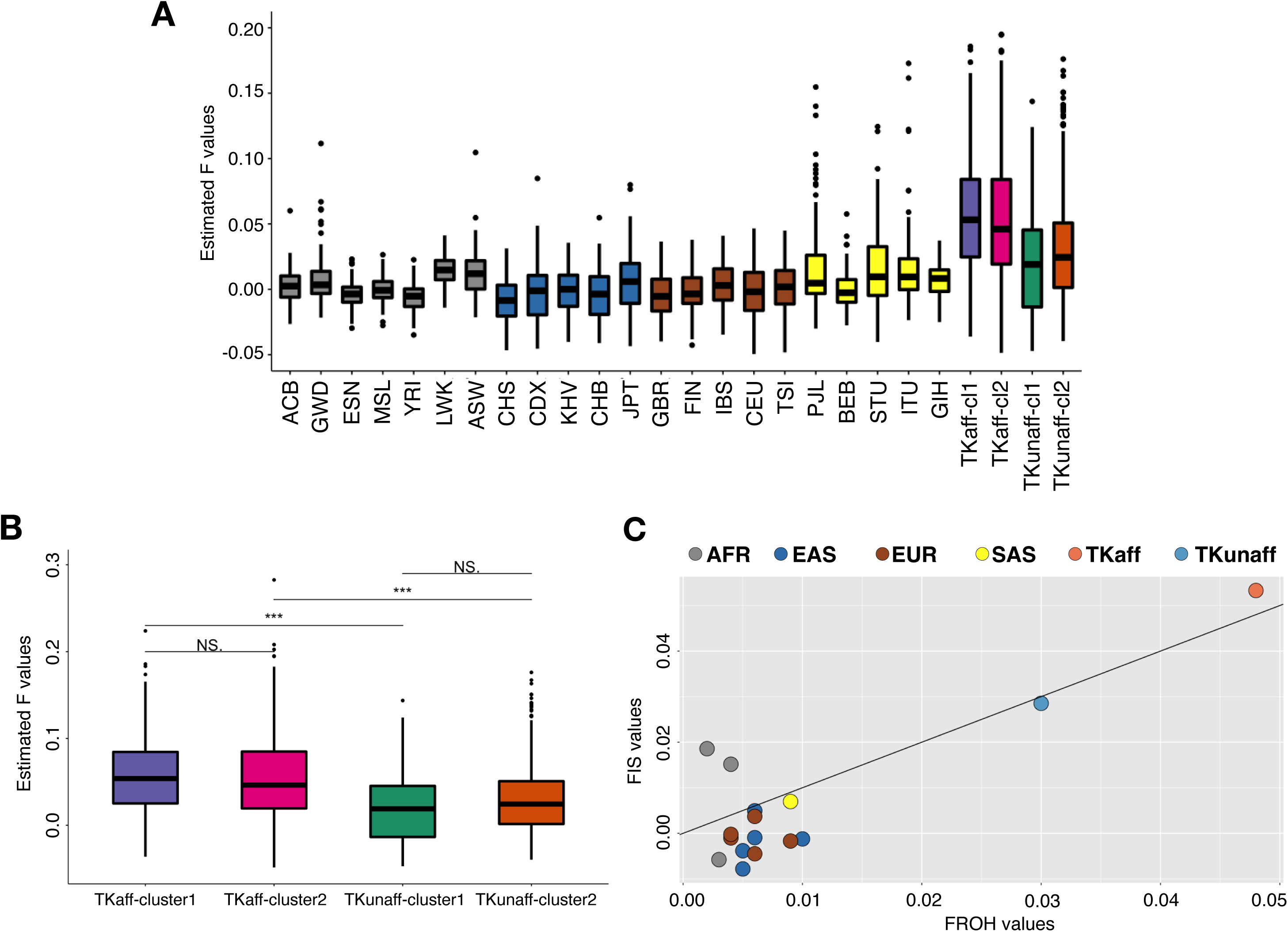
Higher estimated inbreeding coefficient F values in the TK individuals. **A)** The boxplots report the estimated inbreeding coefficient levels (F) calculated from ES data. Both of the TK affected cluster 1 (purple), affected cluster 2 (dark pink) and unaffected cluster 1 (dark green) and unaffected cluster 2 (dark orange) individuals showed higher F values on average compared to the African (AFR) (gray), East Asian (EAS) (dark blue), European (EUR) (brown) and South Asian (SAS) (yellow) subpopulations from the 1000 Genomes project. **B)** The boxplots show the estimated F values of TK affected cluster 1 (TKaff-cluster1) and cluster 2 (TKaff-cluster2) individuals (mean=0.055 vs. 0.052) were significantly higher compared to the TK unaffected cluster 1 (TKunaff-cluster1) and cluster 2 (TKunaff-cluster2) individuals, respectively (Wilcoxon test one-tailed P-values 2.3e-6 and 2.2e-16). There was no significant difference noted between F values of the TK affected cluster 1 and cluster 2 individuals (P-value=0.34) and the TK unaffected cluster 1 and cluster 2 (P=0.11). P-values indicated above each pair of groups compared (*<0.05, **<0.01, ***<0.001, ****<0.0001). Outliers are not shown in the boxplots. **C)** The scatter plot compares the average fraction of individual genome covered by long-sized (> 2.64 Mb) ROHs (FROH) to the average estimated F values (FIS) for the African (AFR), East Asian (EAS), European (EUR), South Asian (SAS) and TK affected (TK-aff) and TK unaffected (TK-unaff) samples.

### Enrichment of long-sized ROH genomic regions in TK cohort individuals

Then we hypothesized that higher F values measured in the TK cohort correlate with an increased length and number of long-sized ROH regions that arose on recently configured “young” haplotypes due to recent parental relatedness. To test this, we identified first ROH regions from ES data using an informatics tool, BafCalculator (https://github.com/BCM-Lupskilab/BafCalculator)^32^ that calculates genomic intervals with absence of heterozygosity, AOH, from unphased ES data as a surrogate measure of ROH. To obtain copy-number neutral ROH regions from the calculated ROH intervals, we excluded a subset of the apparent homozygous regions caused by common variant CNV deletions. Applying Gaussian-mixture modeling with the mclust R package, we classified the genomic intervals for ROH regions detected through BafCalculator into three length classes: short-sized (0.210-0.671 Mb), medium-sized (0.671-2.64 Mb) and long-sized ROHs (>2.64 Mb). To show that those ROH regions in these three different and discrete categories are not overlapping and are formed by different factors (e.g., recent inbreeding and local recombination rate), we examined the distribution of those ROH regions of long-, medium-, and short-sized ROH regions and observed that they were located in different parts of the genome, i.e., they map to different genetic locus intervals, and those regions were distributed non-uniformly across the genome as clearly visualized in the circos plot (Supplementary Figure 5). The non-uniform distribution of ROHs may be formed across the genome because of several physical properties of the human genome: they may involve some genes that are targets of positive selection in a population^57^ or they may include small structural variant (SV) inversions that suppress recombination^58,59^.

We also examined the effect of local recombination rate on the formation of those ROHs of three different and discrete ROH size classes. In alignment with the finding that ROHs of long-sized, medium-sized, and short-sized ROHs distributed in different parts of the genome (Supplementary Figure 5), we found that long-sized ROHs are more likely to contain human genome recombination cold spot regions^60^ compared to the medium-sized (Wilcoxon test one-tailed P-value=1.54e-13) and short-sized ROHs (Wilcoxon test one-tailed P-value=2.08e-29). Overall, our computational analyses showed that we uncovered three different and discrete ROH size classes that do not overlap with each other (distributed in different parts of the genome) and are specific and meaningful for the TK cohort. In addition, our data support the notion that ‘genome geography of recombination rates’ [i.e., positions of ‘coldspots’ versus ‘hotspots’ for recombination] may influence genetic architecture in specific populations.

To investigate the effect of inbreeding level on the genetic architecture of disease traits and to uncover which genomic regions are more contributory to disease phenotypes in the TK population, we compared the ROH length distribution between the TK affected and unaffected participants. As expected, the TK affected participants (with an average of estimated F values=0.053) have a higher level of estimated F values compared to TK unaffected participants (with an average of estimated F values=0.028), given that consanguinity is well established to be a risk factor for rare disorders. This increase in the estimated F levels in TK affected vs. unaffected participants was reflected in an increase in the long-sized ROH total size (median=111.71 Mb vs. 34.41 Mb, Wilcoxon test one-tailed P-value=2.09e-18) and number (median=13 vs. 6, Wilcoxon test one-tailed P-value=6.14e-16), but not medium-sized and short-sized ROHs (Figure 4A and Supplementary Figure 6).

**Figure 4.**
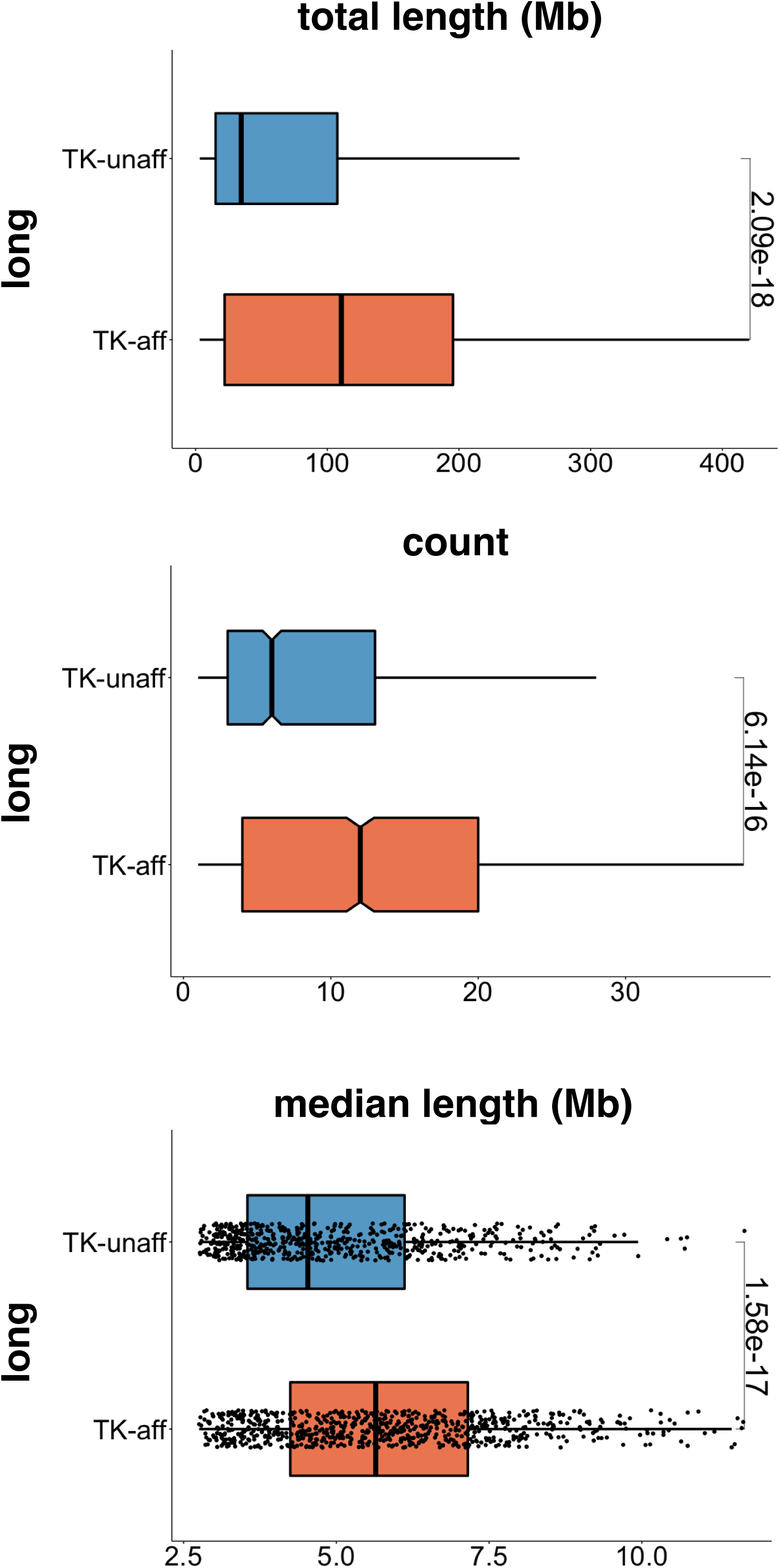
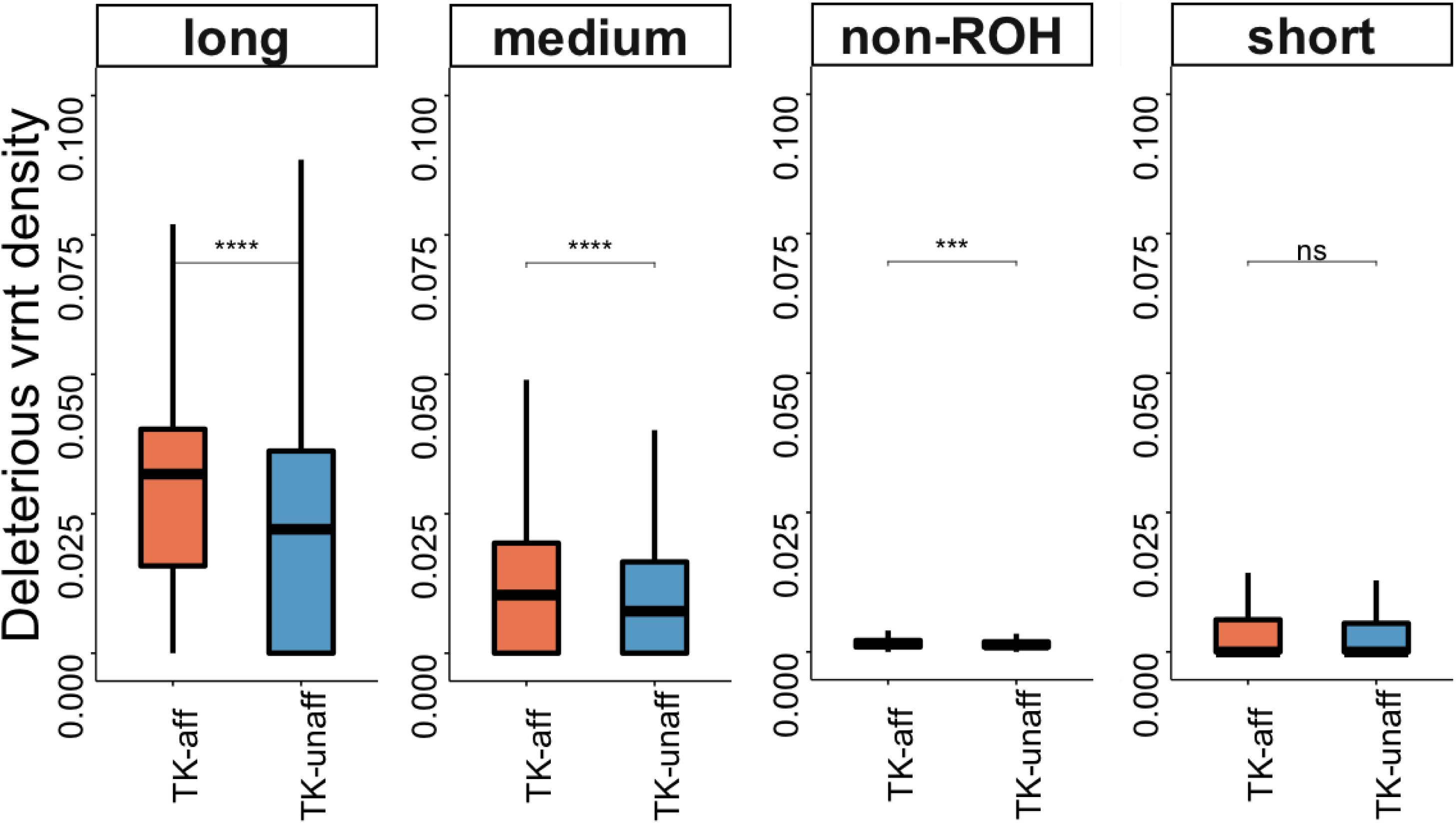
Long-sized ROH regions with increased density of rare deleterious variants are enriched in TK affected participants. **A)** The features of long-sized ROHs were displayed irrespective of their total size (Mb) (top panel), the number of ROH blocks (middle panel) and the median length of ROH blocks (Mb) (bottom panel). In each panel, horizontal boxplots compare the two individual groups as the TK affected (TK-aff) and unaffected (TK-unaff) individuals. A one-sided Wilcoxon rank sum test was used to test the difference in the TK-affected versus TK-unaffected individuals and is indicated to the right of each panel. **B)** We calculated a variant density metric by normalizing the count of rare homozygous deleterious variants to the count of rare homozygous synonymous variants. The density of rare homozygous deleterious variants manifested in long-sized, medium-sized, short-sized ROH regions and not ROH (non-ROH) regions in the TK affected (TK-aff) and TK unaffected (TK-unaff) participant subgroups. We then compared the rare homozygous deleterious variant density between the TK-aff and TK-unaff subgroups in each ROH size group and non-ROHs. P-value significance levels were marked on the top of each pair of groups compared (*<0.05, **<0.01, ***<0.001, ****<0.0001). Outliers were not shown in the boxplots.

In parallel, we performed a correlation analysis to explore the hypothesis that higher estimated F values observed in the TK cohort manifested an increased genome-wide burden of long-sized (>2.64 Mb) ROHs. Our analysis revealed that as the estimated inbreeding levels increase in the TK affected participants, the genome-wide burden of long-sized ROHs increase (ρ=0.83, Supplementary Figure 7A), but not the genome-wide burden of medium-sized ROHs (ρ=0.29, Supplementary Figure 7B) or short-sized ROH segments (ρ=-0.25, Supplementary Figure 7C). This analysis supports the notion that recent inbreeding is most likely to contribute to the formation of long-sized ROHs, but not to the other defined categories (i.e., medium and short). Taken together, both analyses showed that, as the estimated inbreeding levels increase within the TK cohort (TK affected vs. TK unaffected participants and within the TK affected participants), the genome-wide burden of long-sized ROH regions also increases.

We next examined specifically the characteristics of other ROH size categories. The total length of short-sized ROH regions, which result from short homozygous blocks on ancient haplotypes showed a moderate negative correlation with the estimated F values (Supplementary Figure 7C). Concordant with the lower estimated F values in the TK unaffected participants, the short-sized ROH regions in the unaffected participants were greater in total size (median=31 vs. 28.62 Mb, Wilcoxon test one-tailed P-value=2.41e-7) and higher in number (median=126 vs. 122, Wilcoxon test one-tailed P-value=7.65e-8) than those of affected participants (Supplementary Figure 6B). In contrast, medium-sized ROH regions that arise mostly due to background relatedness ^27,57^ were present in a slightly higher number in the TK unaffected compared with those of affected participants (median=34 vs. 32, Wilcoxon test one-tailed P-value=0.04) but did not differ significantly in total length (median=39.11 vs. 39.77 Mb, Wilcoxon test one-tailed P-value=0.199) between the TK affected and unaffected participants (Supplementary Figure 6A).

Taken together, these ROH analyses support the notion that long-, medium-, and short-sized ROH regions result from individual or personal genome population history and dynamics such as consanguinity level or background relatedness ^27,57^ in nuclear families and clans. Of note, the TK affected participants show a significantly increased overall burden of ROH segments per genome compared to unaffected participants, stemming from an increased genome-wide burden of long-sized ROHs.

### Enrichment of predicted deleterious variants in long-sized ROH regions

Then we tested the hypothesis that long-sized ROH regions that arose due to recent parental relatedness would be enriched for rare homozygous deleterious variants, because insufficient generational time would not yet have allowed for selective elimination of such detrimental alleles from a population ^61–63^.We first identified rare homozygous deleterious variants (MAF ≤ 0.05) predicted to be potentially deleterious, by selecting variants above a CADD PHRED-scaled score of >=15 and with a prediction tool algorithm, NMDescPredictor, to predict potential LoF variants^64^. To account for variable mutation rates across different samples and genomic regions, we computed a variant density metric by normalizing the count of rare homozygous deleterious variants to the count of rare homozygous synonymous variants. Since ROH regions are likely to enable deleterious variation to exist in a homozygous form, we then investigated the contribution of ROH regions to deleterious variant density in the TK cohort, by grouping rare homozygous deleterious variants into four groups based on their genomic location. Restated, variants within ROH regions in three different size categories (long-, medium-, and short-sized) or outside of an ROH block (non-ROH) [This is an incomplete sentence]. As expected, in both the TK affected and unaffected participants, we observed an increased level of mutational burden in ROH regions that was most striking in long-sized ROH regions, which are more likely to be shaped by young haplotype blocks including deleterious variation. This was followed by medium-sized ROH, short-sized ROH, and non-ROH regions (Figure 4B).

By comparison, we observed a significantly increased level of rare homozygous deleterious variant density in the TK affected compared to unaffected participants that was most strikingly outlined in long-sized (Wilcoxon test one-tailed P-value= 3.35e-11) and to a lesser extent in medium-sized ROHs (Wilcoxon test one-tailed P-value= 1.9e-9) and non-ROHs (Wilcoxon test one-tailed P-value= 4.36e-2, Figure 4B). Short-sized ROHs do not seem to contribute to the overall rare homozygous deleterious variant density difference observed in ROH regions between the TK affected and unaffected participants (Wilcoxon test one-tailed P-value= 1.75e-1, Figure 4B).

In summary, this genome analysis documented that long-sized ROHs were most enriched for rare homozygous deleterious variations compared to medium-sized, short-sized, and non-ROHs, irrespective of affection status in the TK cohort, indicating a contribution of recent parental relatedness to mutational burden in the TK population (Figure 4B). This further demonstrated that the overall burden of rare homozygous deleterious variation was significantly increased in TK affected compared to unaffected participants and most prominent in long-sized ROHs, conceptually reflecting copy number neutral structural variant haplotypes derived by new mutation and present with low rates of recombination in recent ancestors of the clan.

### Phenotypic and population structure characterization of a mixed disease cohort

The TK participants were recruited into the BHCMG because of suspected Mendelian disorders and presented with a wide variety of clinical phenotypes. To define the range and spectrum of clinical phenotypes, and the genetic traits observed in these individuals, we generated a phenotypic similarity score with the R package ontology Similarity^29,47,48^ and the Resnik approach to information content^65^ to evaluate semantic similarity in a taxonomy between each pair of individuals using the HPO terms^66,67^ recorded in PhenoDB. We performed an unsupervised clustering of those participants based on this phenotypic similarity score, revealing five major ‘disease phenotype groups’ in the TK affected individuals. Clusters 1, 3, and 5 consist mainly of individuals with diverse phenotypic features, which do not fit into a ‘singular general disease category or group’, i.e. higher order HPO term or generalizable disease state or clinical diagnosis. Clusters 2 and 5 are formed of individuals who could be grouped by defined clinical entities largely reflective of complex disease traits, including the hypergonadotropic hypogonadism (HH) cohort^68^ and the neurological disorders cohort^4^, respectively (Figure 5A).

**Figure 5.**
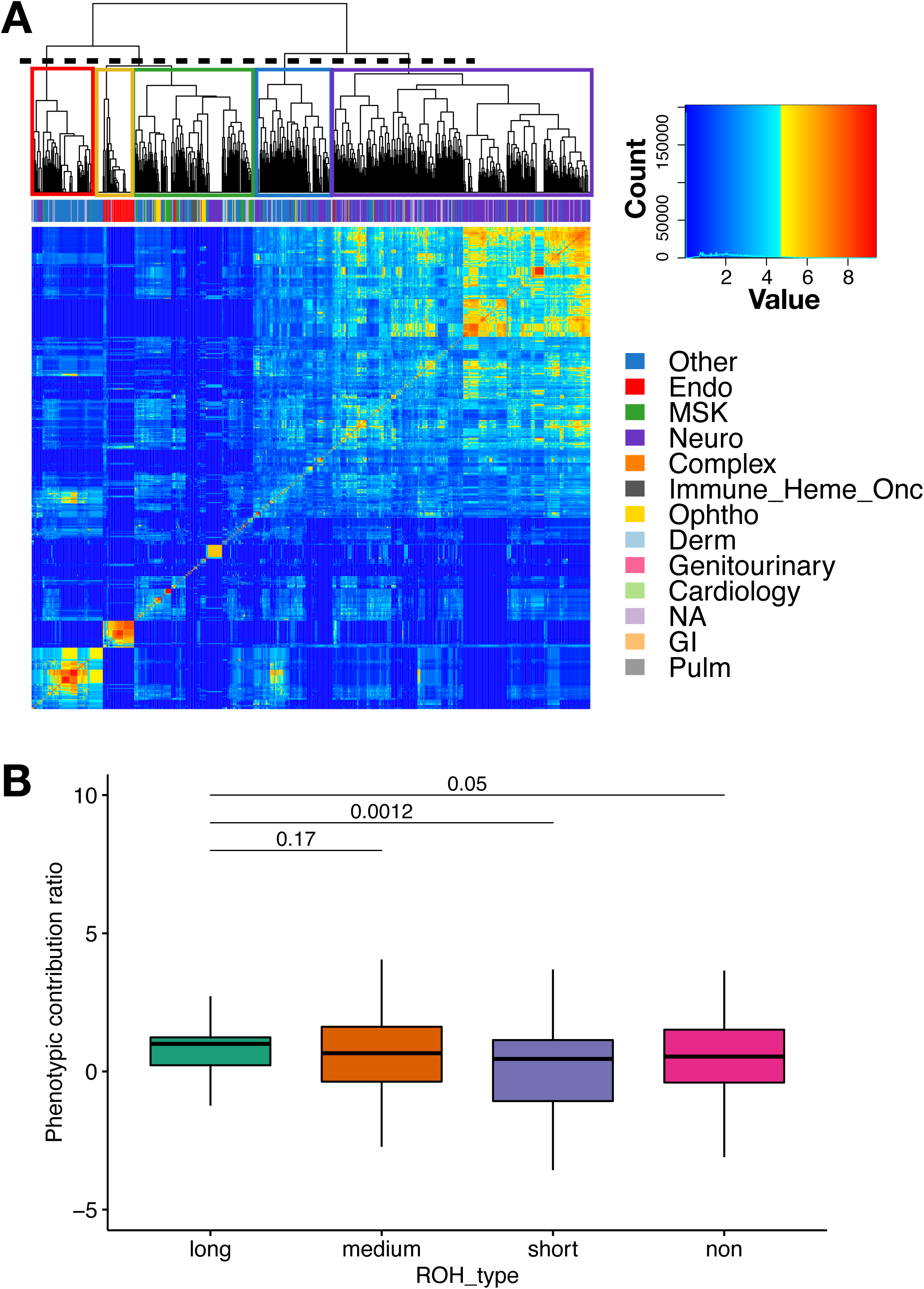
Contribution of variants in ROH to disease trait phenotype. **A)** A heatmap depicting the unsupervised hierarchical clustering of the TK affected patients into five general phenotypic categories or groups by calculating a pairwise phenotypic similarity score (clusters 1 (colored in red), 2 (colored in yellow), 3 (colored in green), 4 (colored in blue) and 5 (colored in purple)). **B)** The comparison between the long-sized ROH group to medium-sized, short-sized ROHs and non-ROHs with regard to the performance of their variants in contribution to the disease trait phenotype presented in the TK patients. The y-axis describes the ratios of z-scores calculated for variants located in a specific ROH size group, e.g., long-sized ROH group, versus z-scores calculated for all variants of interest. Z-scores were calculated for each tested disease gene list performing 1,000 simulations. In each simulation, a permuted disease gene list was selected that has a same number of genes as the tested disease gene list. Then, a similarity score was computed between the patient’s clinical features and the associated HPO terms of that permuted disease gene list.

### Genotype-phenotype analyses revealing ROH-associated genetic architecture of diseases

We then tested the extent to which mutational burden caused by rare homozygous deleterious variants in long-sized ROH regions explains clinical phenotypic features of the individuals in the TK cohort. To this end, we performed an unbiased genotype-phenotype correlation analysis based on the HPO terms^69,70^. In this analysis, we linked the disease genes present with rare coding homozygous and deleterious variation in an individual genome to their related HPO terms for the trait, as defined in the OMIM (https://www.omim.org) clinical synopsis, according to HPO annotation resource databases. Then, for the clinical phenotypic features of each patient recorded in PhenoDB^54^, we compared the associated HPO term sets to the merged HPO term sets for the disease genes defined for each ROH category, using a semantic similarity score metric that controls for the number of genes and ROH block size from a permutation approach^29,47,48^. These analyses showed that, in TK affected individuals, genes with rare homozygous deleterious variants in long-sized ROH regions are most informative for clinical phenotypic features objectively assessed and compared, fromthe information submitted to and collated within PhenoDB^54^ compared to medium-sized ROHs (P-value=1.7e-1), short-sized ROHs (P-value=1.2e-3) and non-ROHs (P-value=5e-2) (Figure 5B).

These analyses also revealed that the top-ranking genes with rare coding homozygous deleterious variant alleles were located in the long-sized ROH regions in 152 patients (phenotypic similarity score >=1). Importantly, 75 out of those 152 patients were found to carry rare coding homozygous and deleterious variation located in >=2 OMIM disease genes that contribute significantly to the patients’ phenotypes on the basis of phenotypic similarity score (score>=1). In line with these findings, we indicate a subset of the TK affected participants in the cohort present with neurodevelopmental disorders (NDD; N=234) that were analyzed by expert clinicians. In 176 of those 234 studied participants (75.2%), a plausible and genetically parsimonious molecular etiology due to rare coding variation was identified. Importantly, out of those 176 participants, 51 families (51/176 = 28.9%) were identified with multi-locus pathogenic variation (MPV), mostly driven by ROHs^7^. The retrospective analysis of those 176 participants revealed that cases with diagnoses involving >=2 disease gene loci (N=51) were found to have significantly higher F values (Wilcoxon test one-tailed P-value= 9.35e-5, Figure 6A) and significantly increased total span of long-sized ROHs (Wilcoxon test one-tailed P-value=1.06e-4, Figure 6B) compared to cases with diagnoses involving one (1) disease gene locus. On the other hand, we found that there were no significant differences between these two groups of patients in terms of total span of medium-sized ROHs (Wilcoxon test one-tailed P-value=0.223, Figure 6C). In summary, our results provide compelling evidence that an increased level of consanguinity observed in the TK affected compared to unaffected participants is correlated with a greater total span of long-sized ROH blocks and significantly increased level of rare homozygous deleterious variant density, thereby an elevated mutational burden that likely contributes to the disease trait(s) phenotype observed in the individuals studied.

**Figure 6.**
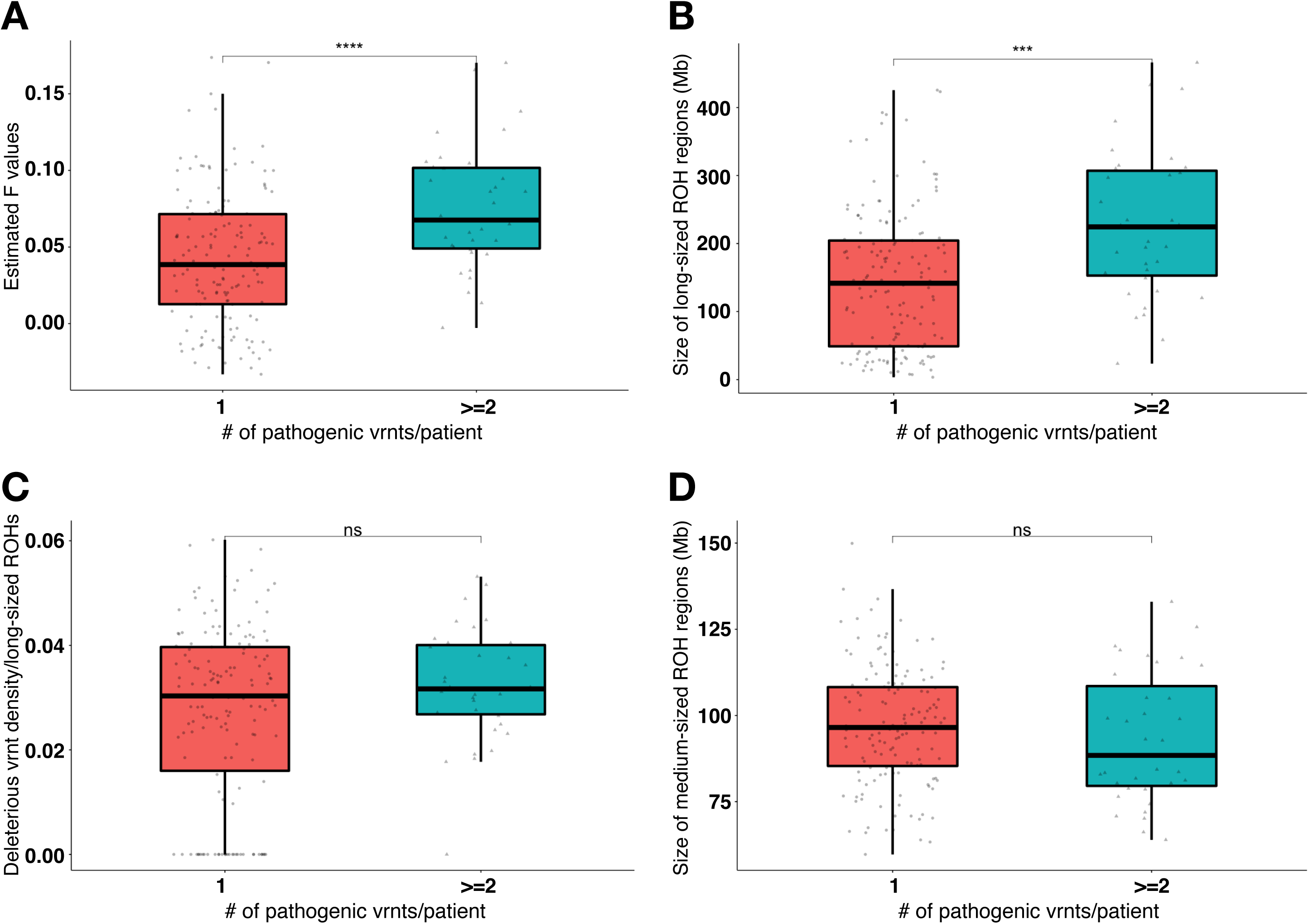
An elevated mutational burden in long-sized ROHs can produce blended phenotypes. A retrospective analysis of previously published cases (N=217) compared the cases with diagnoses involving >=2 disease gene loci (N=40) compared to cases with diagnoses involving 1 disease gene locus in terms of **A)** estimated F values, **B)** total span of long-sized ROHs, **C)** deleterious variant density manifested in long-sized ROHs and **D)** total span of medium-sized ROHs.

## DISCUSSION

Our results provide compelling evidence that an increased level of consanguinity observed in the TK affected compared to unaffected participants is correlated with a greater total span of long-sized ROH blocks and significantly increased level of rare homozygous deleterious variant density, thereby an elevated mutational burden that likely contributes to rare disease trait(s). Comprehensive catalogues of common and rare human genetic variation stored in population variant databases, such as ExAC^71^, gnomAD^55^ (https://gnomad.broadinstitute.org), and ARIC^64,72^, have proven to be powerful resources and interpretive tools to identify rare, ultra-rare, and potentially pathogenic variations that influence gene action and expression of Mendelian disease traits. However, many world populations remain for which genetic variation is underrepresented or entirely absent in such databases, including populations with more prevalent consanguinity and/or a considerable amount of admixture, such as the TK population ^41^.

Beyond the elucidation of the biological basis and molecular pathogenesis of disease, the mapping of a locus at which variation might contribute to disease analysis and characterization of molecular features in personal genomes can potentially provide some insights into the genetic architecture contributing to disease traits in a population and evolution of personal genomes. From our TK population cohort, we show that investigation of the molecular features of ROH regions culled from personal unphased ES data through BafCalculator (https://github.com/BCM-Lupskilab/BafCalculator) ^32^ and classified through Gaussian-mixture machine learning modeling into three length ROH classes specific to the TK population can provide insights into the haplotype derivation and genetic architecture of disease. Long-sized ROH (>2.64 Mb) regions that arose on recently configured haplotype blocks were greater in size (both total and median length) and number in the genomes of the TK affected compared to TK unaffected participants (median=111.71 Mb vs. 34.41 Mb, Wilcoxon test one-tailed P-value=2.09e-18 and median=13 vs. 6, Wilcoxon test one-tailed P-value=6.14e-16) without any significant cluster-specific differences (Figure 4A, Supplementary Figures 8 and 9). This finding is concordant with higher estimated F values (0.053 vs. 0.028, Wilcoxon test one-tailed P-value= 2.35e^-18^) and high level of consanguinity due to recent shared ancestors in the TK population ^56^. Characterization of ROH genomic intervals also revealed that long-sized ROH regions on newly derived haplotypes are enriched particularly with rare homozygous deleterious variants specifically in the TK affected compared to TK unaffected participants (Wilcoxon test one-tailed P-value= 3.35e-11). These findings support the notion that the inbreeding level results in an increase in the genome-wide burden of long-sized ROH along with an increase in their rare homozygous deleterious variation density (Figure 4A and B, and Supplementary Figure 10).

To test the contribution of those ROH-derived rare variant combinations to Mendelian rare disease traits, we performed a large-scale analysis of genotype-phenotype correlations in the TK cohort from their clinical features captured as structured HPO terms^66,67^ (https://hpo.jax.org/app/download/annotation) recorded in PhenoDB^54^. A systematic, integrative, quantitative analysis revealed that the combinatorial phenotypic effect of ultra-rare variants embedded within long-size ROH regions strongly explain the observed rare disease trait(s) in the TK cohort. Corroborating these findings, a retrospective analysis of previously published cases (N=176)^3,4,7,73^ demonstrated that the cases with multi-locus pathogenic variation (MPV; N=51) (diagnoses involving >=2 disease gene loci) are more likely to have an increased genome-wide burden of long-sized ROH regions (Wilcoxon test one-tailed P-value=1.06e-4, Figure 6B), corresponding to higher estimated F values (Wilcoxon test one-tailed P-value= 9.35e-5, Figure 6A) compared to cases with diagnoses involving one disease gene locus. That these variants were ultra-rare within the TK population itself suggests that they may represent new mutation events within individual families or the clan, the most basic unit of a population. Population substructure ultimately driving formation of these newly configured – and unique – disease haplotypes can be rapidly brought to homozygosity through IBD: each shaped by recombination, characterized by new mutation, and even more rare than their constituent alleles^74^.

This phenomenon of new mutations as ultra-rare variants on a newly derived haplotype may provide important insights into the molecular etiology of disease traits, and further amplify several prior observations including: (1) the role of multilocus variation underlying some cases of apparent phenotypic expansion^3,29,32,47^; (2) homozygosity of ultra-rare pathogenic variation in *PRUNE* (HGNC:13420) in sect-driven population isolates^4,75,76^ in stark contrast with the contribution of rare (but not ultra-rare) *CLP1* (HGNC:16999) founder alleles to microcephaly and neurodevelopmental disease^77,78^; (3) the seemingly now-common identification of genes for which biallelic variation can lead to disease traits previously categorized as strictly ‘dominant Mendelian loci’-traits^7,79–81^, and (4) the emergence of a founder allele arisen by identity-by-state (IBS) in Steel syndrome of the geographically isolated population of the Commonwealth of Puerto Rico versus clan genomics derived *COL27A1* (HGNC:22986) pathogenic variation in the TK population^37^.

Our findings also highlight the advantage of merging per-locus variation with genomics (inherited vs. *de novo* mutations) for gleaning insights into the genetic architecture contributing to disease in a population. Multiple genetic changes could be brought together per locus to generate unique haplotype blocks. One example of this phenomenon is the observation of uniparental isodisomy (UPD) manifest by a long tract of homozygosity on the entirety of chromosome 7 explaining both the short stature phenotype in addition to the expected clinical features of cystic fibrosis (CF, OMIM #219700) in a child^51^. Viewed from such a perspective, disease phenotypes resulting from UPD, and epigenetic/imprinting diseases may benefit from haplotype phased genomes and long read technologies that differentiate methylated W-C bases^82,83^. Another example of per-locus genetic studies is the modulation of disease risk through gene expression and dosage effects of regulatory common variant (expression quantitative trait loci, eQTLs) haplotype configurations of coding pathogenic variants and CNVs^5,84–88^. The high mutation rate of recurrent genomic deletions [e.g., driven by a special type of mutational mechanism, nonallelic homologous recombination (NAHR)] may make these loci a significant contributor to AR disease trait loci in populations due to a compound heterozygous CNV + SNV allelic combination^89^.

To apply a ‘merging’ of genetics (per locus variation) and genomics (inherited ‘variome’ and *de novo* mutations) thinking to the clinic, our data emphasize identifying and systematically analyzing the ROH genomic intervals culled from the patient’s personal unphased ES data with BafCalculator (https://github.com/BCM-Lupskilab/BafCalculator)^32^. If the degree of parental relatedness as judged by the ROH size culled from unphased ES data suggests that the patient may come from a clan with a high estimated coefficient of consanguinity, rare homozygous variants mapping within the ROH regions and their combinatorial effect could be prioritized in molecular analyses^3,29,32,47^. Culling ROH regions from unphased clinical ES data with BafCalculator (https://github.com/BCM-Lupskilab/BafCalculator)^32^ may also enhance the discovery of pathogenic homozygous or hemizygous exonic CNV deletions arising on newly derived structural variant (SV) haplotypes in a clan homozygosed by IBD ^90–93^.

In summary, the findings in this study support the Clan Genomics hypothesis^94–96^, which suggests that newly configured haplotypes resulting from recent mutations play a role in Mendelian diseases. The rapid generation of disease haplotypes driven by population substructure and shaped by recombination contributes to the occurrence of these rare disease-causing haplotypes. That these haplotypes are even more rare than their constituent alleles reinforces the importance of newly arisen mutations in rare recessive disease traits.

## DATA and CODE AVAILABILITY

All variants reported herein have been aggregated within a TK variome database that is publicly available as a research and molecular diagnostic community resource (https://turkishvariomedb.shinyapps.io/tvdb/). The code generated during this study is available at (https://github.com/BCM-Lupskilab/BafCalculator). Exome variant data have been deposited to dbGaP (study accession phs000711.v5.p1) and/or AnVIL for all cases for which written, informed consent for sharing of data through controlled-access databases has been obtained, in keeping with our IRB and BCM protocol H-29697. Requests for further information on raw data, genomic and phenotypic analyses, DNA samples may be directed to, and will be fulfilled by Lead Contact James R. Lupski.

## ACKNOWLEDGMENTS

We thank all patients, their families, and their referring physicians who submitted samples for genomic studies. No additional compensation was offered for these studies.

## AUTHOR CONTRIBUTIONS

Conceptualization: Z.C.A., X.S., F.C., D.P. and J.R.L.; Data curation: Z.C.A., X.S., F.C., T.G. and T.B.Y.; Formal Analysis: Z.C.A., X.S., F.C., D.P. and J.R.L; Supervision: J.R.L. and D.V.; Project administration: S.N.J., T.G., C.M.B.C., D.M.M., P.L, E.B., R.A.G., C.A.S. and J.E.P.; Funding acquisition: R.A.G, J.E.P., D.V. and J.R.L.; Clinical resources: D.P., E.K., Y.B., T.M., A.H, V.R.S, R.A.L. and N.S.; Writing original draft: Z.C.A., X.S., F.C., D.P. and J.R.L; Writing, review, and editing: Z.C.A., X.S., F.C., D.P., R.A.L., C.A.S, C.M.B.C, J.E.P and J.R.L. All contributing co-authors have read, edited, and approved the final manuscript.

## ETHICS DECLARATION

This study was approved by the Institutional Review Board at Baylor College of Medicine (IRB protocol # H-29697). We recruited 1,416 unrelated TK individuals (669 females and 747 males) in the Baylor Hopkins Center for Mendelian Genomics (BHCMG) cohort (data freeze: December 2011- October 2020) after all relevant subjects or legally authorized representatives provided written informed consent for the use of their DNA and personal genomes for the identification of potential disease-contributing variants and for broad data sharing.

## WEB RESOURCES

1000 Genomes Project Database, http://ftp.1000genomes.ebi.ac.uk/vol1/ftp/release/20130502/

Atherosclerosis Risk in Communities Study (ARIC) Database, http://www2.cscc.unc.edu/aric/

BafCalculator ROH Detection Tool, https://github.com/BCM-Lupskilab/BafCalculator

dbGaP, https://www.ncbi.nlm.nih.gov/gap

ExAC Browser, http://exac.broadinstitute.org/

gnomAD Browser, https://gnomad.broadinstitute.org/

Human Genome Diversity Panel, https://www.internationalgenome.org/data-portal/data-collection/hgdp

Human Phenotype Ontology (HPO) Terms Annotation, https://hpo.jax.org/app/download/annotation

Mercury Analysis Pipeline, https://www.hgsc.bcm.edu/software/mercury

NMDescPredictor, https://nmdprediction.shinyapps.io/nmdescpredictor/

NMD Escape Intolerance Score, https://nmdprediction.shinyapps.io/nmdescintolerancescore/

OMIM, http://www.omim.org/

PhenoDB, https://phenodb.org/

The Greater Middle Eastern Variome Database, http://igm.ucsd.edu/gme/

Turkish Variome Database, https://turkishvariomedb.shinyapps.io/tvdb/

## References

1. Reich, D.E., and Lander, E.S. (2001). On the allelic spectrum of human disease. Trends Genet 17, 502–510. 10.1016/s0168-9525(01)02410-6.

2. Wang, W.Y., Barratt, B.J., Clayton, D.G., and Todd, J.A. (2005). Genome-wide association studies: theoretical and practical concerns. Nat Rev Genet 6, 109–118. 10.1038/nrg1522.

3. Pehlivan, D., Bayram, Y., Gunes, N., Coban Akdemir, Z., Shukla, A., Bierhals, T., Tabakci, B., Sahin, Y., Gezdirici, A., Fatih, J.M., et al. (2019). The Genomics of Arthrogryposis, a Complex Trait: Candidate Genes and Further Evidence for Oligogenic Inheritance. Am J Hum Genet 105, 132–150. 10.1016/j.ajhg.2019.05.015.

4. Karaca, E., Harel, T., Pehlivan, D., Jhangiani, S.N., Gambin, T., Coban Akdemir, Z., Gonzaga-Jauregui, C., Erdin, S., Bayram, Y., Campbell, I.M., et al. (2015). Genes that Affect Brain Structure and Function Identified by Rare Variant Analyses of Mendelian Neurologic Disease. Neuron 88, 499–513. 10.1016/j.neuron.2015.09.048.

5. Wu, N., Ming, X., Xiao, J., Wu, Z., Chen, X., Shinawi, M., Shen, Y., Yu, G., Liu, J., Xie, H., et al. (2015). TBX6 null variants and a common hypomorphic allele in congenital scoliosis. N Engl J Med 372, 341–350. 10.1056/NEJMoa1406829.

6. Groopman, E.E., Povysil, G., Goldstein, D.B., and Gharavi, A.G. (2020). Rare genetic causes of complex kidney and urological diseases. Nat Rev Nephrol 16, 641–656. 10.1038/s41581-020-0325-2.

7. Mitani, T., Isikay, S., Gezdirici, A., Gulec, E.Y., Punetha, J., Fatih, J.M., Herman, I., Akay, G., Du, H., Calame, D.G., et al. (2021). High prevalence of multilocus pathogenic variation in neurodevelopmental disorders in the Turkish population. Am J Hum Genet 108, 1981–2005. 10.1016/j.ajhg.2021.08.009.

8. Alkuraya, F.S. (2010). Homozygosity mapping: one more tool in the clinical geneticist’s toolbox. Genet Med 12, 236–239. 10.1097/GIM.0b013e3181ceb95d.

9. Alkuraya, F.S. (2010). Autozygome decoded. Genet Med 12, 765–771. 10.1097/GIM.0b013e3181fbfcc4.

10. Alkuraya, F.S. (2021). How the human genome transformed study of rare diseases. Nature 590, 218–219. 10.1038/d41586-021-00294-7.

11. Torkamani, A., Pham, P., Libiger, O., Bansal, V., Zhang, G., Scott-Van Zeeland, A.A., Tewhey, R., Topol, E.J., and Schork, N.J. (2012). Clinical implications of human population differences in genome-wide rates of functional genotypes. Front Genet 3, 211. 10.3389/fgene.2012.00211.

12. Tennessen, J.A., Bigham, A.W., O’Connor, T.D., Fu, W., Kenny, E.E., Gravel, S., McGee, S., Do, R., Liu, X., Jun, G., et al. (2012). Evolution and functional impact of rare coding variation from deep sequencing of human exomes. Science 337, 64–69. 10.1126/science.1219240.

13. Lohmueller, K.E., Indap, A.R., Schmidt, S., Boyko, A.R., Hernandez, R.D., Hubisz, M.J., Sninsky, J.J., White, T.J., Sunyaev, S.R., Nielsen, R., et al. (2008). Proportionally more deleterious genetic variation in European than in African populations. Nature 451, 994–997. 10.1038/nature06611.

14. Clark, D.W., Okada, Y., Moore, K.H.S., Mason, D., Pirastu, N., Gandin, I., Mattsson, H., Barnes, C.L.K., Lin, K., Zhao, J.H., et al. (2019). Associations of autozygosity with a broad range of human phenotypes. Nat Commun 10, 4957. 10.1038/s41467-019-12283-6.

15. Joshi, P.K., Esko, T., Mattsson, H., Eklund, N., Gandin, I., Nutile, T., Jackson, A.U., Schurmann, C., Smith, A.V., Zhang, W., et al. (2015). Directional dominance on stature and cognition in diverse human populations. Nature 523, 459–462. 10.1038/nature14618.

16. Keller, M.C., Simonson, M.A., Ripke, S., Neale, B.M., Gejman, P.V., Howrigan, D.P., Lee, S.H., Lencz, T., Levinson, D.F., Sullivan, P.F., and Schizophrenia Psychiatric Genome-Wide Association Study, C. (2012). Runs of homozygosity implicate autozygosity as a schizophrenia risk factor. PLoS Genet 8, e1002656. 10.1371/journal.pgen.1002656.

17. Yengo, L., Wray, N.R., and Visscher, P.M. (2019). Extreme inbreeding in a European ancestry sample from the contemporary UK population. Nat Commun 10, 3719. 10.1038/s41467-019-11724-6.

18. Smith, C.A.B. (1953). The Detection of Linkage in Human Genetics. Journal of the Royal Statistical Society. Series B (Methodological) 15, 153–192.

19. Morton, N.E. (1991). Genetic epidemiology of hearing impairment. Ann N Y Acad Sci 630, 16–31.

20. Lander, E.S., and Botstein, D. (1987). Homozygosity mapping: a way to map human recessive traits with the DNA of inbred children. Science 236, 1567–1570.

21. Broman, K.W., and Weber, J.L. (1999). Long homozygous chromosomal segments in reference families from the centre d’Etude du polymorphisme humain. Am J Hum Genet 65, 1493–1500. 10.1086/302661.

22. Seelow, D., Schuelke, M., Hildebrandt, F., and Nurnberg, P. (2009). HomozygosityMapper--an interactive approach to homozygosity mapping. Nucleic Acids Res 37, W593–599. 10.1093/nar/gkp369.

23. Houwen, R.H., Baharloo, S., Blankenship, K., Raeymaekers, P., Juyn, J., Sandkuijl, L.A., and Freimer, N.B. (1994). Genome screening by searching for shared segments: mapping a gene for benign recurrent intrahepatic cholestasis. Nat Genet 8, 380–386. 10.1038/ng1294-380.

24. Alazami, A.M., Patel, N., Shamseldin, H.E., Anazi, S., Al-Dosari, M.S., Alzahrani, F., Hijazi, H., Alshammari, M., Aldahmesh, M.A., Salih, M.A., et al. (2015). Accelerating novel candidate gene discovery in neurogenetic disorders via whole-exome sequencing of prescreened multiplex consanguineous families. Cell Rep 10, 148–161. 10.1016/j.celrep.2014.12.015.

25. Alazami, A.M., Shaheen, R., Alzahrani, F., Snape, K., Saggar, A., Brinkmann, B., Bavi, P., Al-Gazali, L.I., and Alkuraya, F.S. (2009). *FREM1* mutations cause bifid nose, renal agenesis, and anorectal malformations syndrome. Am J Hum Genet 85, 414–418. 10.1016/j.ajhg.2009.08.010.

26. Alazami, A.M., Al-Saif, A., Al-Semari, A., Bohlega, S., Zlitni, S., Alzahrani, F., Bavi, P., Kaya, N., Colak, D., Khalak, H., et al. (2008). Mutations in *C2orf37*, encoding a nucleolar protein, cause hypogonadism, alopecia, diabetes mellitus, mental retardation, and extrapyramidal syndrome. Am J Hum Genet 83, 684–691. 10.1016/j.ajhg.2008.10.018.

27. McQuillan, R., Leutenegger, A.L., Abdel-Rahman, R., Franklin, C.S., Pericic, M., Barac-Lauc, L., Smolej-Narancic, N., Janicijevic, B., Polasek, O., Tenesa, A., et al. (2008). Runs of homozygosity in European populations. Am J Hum Genet 83, 359–372. 10.1016/j.ajhg.2008.08.007.

28. Kirin, M., McQuillan, R., Franklin, C.S., Campbell, H., McKeigue, P.M., and Wilson, J.F. (2010). Genomic runs of homozygosity record population history and consanguinity. PLoS One 5, e13996. 10.1371/journal.pone.0013996.

29. Posey, J.E., Harel, T., Liu, P., Rosenfeld, J.A., James, R.A., Coban Akdemir, Z.H., Walkiewicz, M., Bi, W., Xiao, R., Ding, Y., et al. (2017). Resolution of Disease Phenotypes Resulting from Multilocus Genomic Variation. N Engl J Med 376, 21–31. 10.1056/NEJMoa1516767.

30. Kaiser, V.B., Svinti, V., Prendergast, J.G., Chau, Y.Y., Campbell, A., Patarcic, I., Barroso, I., Joshi, P.K., Hastie, N.D., Miljkovic, A., et al. (2015). Homozygous loss-of-function variants in European cosmopolitan and isolate populations. Hum Mol Genet 24, 5464–5474. 10.1093/hmg/ddv272.

31. Ceballos, F.C., Joshi, P.K., Clark, D.W., Ramsay, M., and Wilson, J.F. (2018). Runs of homozygosity: windows into population history and trait architecture. Nat Rev Genet 19, 220–234. 10.1038/nrg.2017.109.

32. Karaca, E., Posey, J.E., Coban Akdemir, Z., Pehlivan, D., Harel, T., Jhangiani, S.N., Bayram, Y., Song, X., Bahrambeigi, V., Yuregir, O.O., et al. (2018). Phenotypic expansion illuminates multilocus pathogenic variation. Genet Med 20, 1528–1537. 10.1038/gim.2018.33.

33. Katsanis, N., Ansley, S.J., Badano, J.L., Eichers, E.R., Lewis, R.A., Hoskins, B.E., Scambler, P.J., Davidson, W.S., Beales, P.L., and Lupski, J.R. (2001). Triallelic inheritance in Bardet-Biedl syndrome, a Mendelian recessive disorder. Science 293, 2256–2259. 10.1126/science.1063525.

34. Bejjani, B.A., Lewis, R.A., Tomey, K.F., Anderson, K.L., Dueker, D.K., Jabak, M., Astle, W.F., Otterud, B., Leppert, M., and Lupski, J.R. (1998). Mutations in *CYP1B1*, the gene for cytochrome P4501B1, are the predominant cause of primary congenital glaucoma in Saudi Arabia. Am J Hum Genet 62, 325–333. 10.1086/301725.

35. Bejjani, B.A., Stockton, D.W., Lewis, R.A., Tomey, K.F., Dueker, D.K., Jabak, M., Astle, W.F., and Lupski, J.R. (2000). Multiple *CYP1B1* mutations and incomplete penetrance in an inbred population segregating primary congenital glaucoma suggest frequent de novo events and a dominant modifier locus. Hum Mol Genet 9, 367–374. 10.1093/hmg/9.3.367.

36. Gonzaga-Jauregui, C., Harel, T., Gambin, T., Kousi, M., Griffin, L.B., Francescatto, L., Ozes, B., Karaca, E., Jhangiani, S.N., Bainbridge, M.N., et al. (2015). Exome Sequence Analysis Suggests that Genetic Burden Contributes to Phenotypic Variability and Complex Neuropathy. Cell Rep 12, 1169–1183. 10.1016/j.celrep.2015.07.023.

37. Gonzaga-Jauregui, C., Yesil, G., Nistala, H., Gezdirici, A., Bayram, Y., Nannuru, K.C., Pehlivan, D., Yuan, B., Jimenez, J., Sahin, Y., et al. (2020). Functional biology of the Steel syndrome founder allele and evidence for clan genomics derivation of *COL27A1* pathogenic alleles worldwide. Eur J Hum Genet 28, 1243–1264. 10.1038/s41431-020-0632-x.

38. Hashmi, M.A. (1997). Frequency of consanguinity and its effect on congenital malformation--a hospital based study. J Pak Med Assoc 47, 75–78.

39. Bittles, A.H., and Black, M.L. (2010). Evolution in health and medicine Sackler colloquium: Consanguinity, human evolution, and complex diseases. Proc Natl Acad Sci U S A 107 Suppl 1, 1779–1786. 10.1073/pnas.0906079106.

40. Bittles, A. (2001). Consanguinity and its relevance to clinical genetics. Clin Genet 60, 89–98.

41. Tuncbilek, E., and Koc, I. (1994). Consanguineous marriage in Turkey and its impact on fertility and mortality. Ann Hum Genet 58, 321–329.

42. Reid, J.G., Carroll, A., Veeraraghavan, N., Dahdouli, M., Sundquist, A., English, A., Bainbridge, M., White, S., Salerno, W., Buhay, C., et al. (2014). Launching genomics into the cloud: deployment of Mercury, a next generation sequence analysis pipeline. BMC Bioinformatics 15, 30. 10.1186/1471-2105-15-30.

43. Challis, D., Yu, J., Evani, U.S., Jackson, A.R., Paithankar, S., Coarfa, C., Milosavljevic, A., Gibbs, R.A., and Yu, F. (2012). An integrative variant analysis suite for whole exome next-generation sequencing data. BMC Bioinformatics 13, 8. 10.1186/1471-2105-13-8.

44. Farek, J., Hughes, D., Mansfield, A., Krasheninina, O., Nasser, W., Sedlazeck, F.J., Khan, Z., Venner, E., Metcalf, G., Boerwinkle, E., et al. (2018). xAtlas: Scalable small variant calling across heterogeneous next-generation sequencing experiments. BioRxiv.

45. Bainbridge, M.N., Wiszniewski, W., Murdock, D.R., Friedman, J., Gonzaga-Jauregui, C., Newsham, I., Reid, J.G., Fink, J.K., Morgan, M.B., Gingras, M.C., et al. (2011). Whole-genome sequencing for optimized patient management. Sci Transl Med 3, 87re83. 10.1126/scitranslmed.3002243.

46. Wang, K., Li, M., and Hakonarson, H. (2010). ANNOVAR: functional annotation of genetic variants from high-throughput sequencing data. Nucleic Acids Res 38, e164. 10.1093/nar/gkq603.

47. Liu, P., Meng, L., Normand, E.A., Xia, F., Song, X., Ghazi, A., Rosenfeld, J., Magoulas, P.L., Braxton, A., Ward, P., et al. (2019). Reanalysis of Clinical Exome Sequencing Data. N Engl J Med 380, 2478–2480. 10.1056/NEJMc1812033.

48. James, R.A., Campbell, I.M., Chen, E.S., Boone, P.M., Rao, M.A., Bainbridge, M.N., Lupski, J.R., Yang, Y., Eng, C.M., Posey, J.E., and Shaw, C.A. (2016). A visual and curatorial approach to clinical variant prioritization and disease gene discovery in genome-wide diagnostics. Genome Med 8, 13. 10.1186/s13073-016-0261-8.

49. Olshen, A.B., Venkatraman, E.S., Lucito, R., and Wigler, M. (2004). Circular binary segmentation for the analysis of array-based DNA copy number data. Biostatistics 5, 557–572. 10.1093/biostatistics/kxh008.

50. Huber, W., Carey, V.J., Gentleman, R., Anders, S., Carlson, M., Carvalho, B.S., Bravo, H.C., Davis, S., Gatto, L., Girke, T., et al. (2015). Orchestrating high-throughput genomic analysis with Bioconductor. Nat Methods 12, 115–121. 10.1038/nmeth.3252.

51. Spence, J.E., Perciaccante, R.G., Greig, G.M., Willard, H.F., Ledbetter, D.H., Hejtmancik, J.F., Pollack, M.S., O’Brien, W.E., and Beaudet, A.L. (1988). Uniparental disomy as a mechanism for human genetic disease. Am J Hum Genet 42, 217–226.

52. Fromer, M., and Purcell, S.M. (2014). Using XHMM Software to Detect Copy Number Variation in Whole-Exome Sequencing Data. Curr Protoc Hum Genet 81, 7 23 21-21. 10.1002/0471142905.hg0723s81.

53. Quinlan, A.R. (2014). BEDTools: The Swiss-Army Tool for Genome Feature Analysis. Curr Protoc Bioinformatics 47, 11 12 11-34. 10.1002/0471250953.bi1112s47.

54. Sobreira, N., Schiettecatte, F., Boehm, C., Valle, D., and Hamosh, A. (2015). New tools for Mendelian disease gene identification: PhenoDB variant analysis module; and GeneMatcher, a web-based tool for linking investigators with an interest in the same gene. Hum Mutat 36, 425–431. 10.1002/humu.22769.

55. Karczewski, K.J., Francioli, L.C., Tiao, G., Cummings, B.B., Alfoldi, J., Wang, Q., Collins, R.L., Laricchia, K.M., Ganna, A., Birnbaum, D.P., et al. (2020). The mutational constraint spectrum quantified from variation in 141,456 humans. Nature 581, 434–443. 10.1038/s41586-020-2308-7.

56. Scott, E.M., Halees, A., Itan, Y., Spencer, E.G., He, Y., Azab, M.A., Gabriel, S.B., Belkadi, A., Boisson, B., Abel, L., et al. (2016). Characterization of Greater Middle Eastern genetic variation for enhanced disease gene discovery. Nat Genet 48, 1071–1076. 10.1038/ng.3592.

57. Pemberton, T.J., Absher, D., Feldman, M.W., Myers, R.M., Rosenberg, N.A., and Li, J.Z. (2012). Genomic patterns of homozygosity in worldwide human populations. Am J Hum Genet 91, 275–292. 10.1016/j.ajhg.2012.06.014.

58. Curtis, D., Vine, A.E., and Knight, J. (2008). Study of regions of extended homozygosity provides a powerful method to explore haplotype structure of human populations. Ann Hum Genet 72, 261–278. 10.1111/j.1469-1809.2007.00411.x.

59. Porubsky, D., Hops, W., Ashraf, H., Hsieh, P., Rodriguez-Martin, B., Yilmaz, F., Ebler, J., Hallast, P., Maria Maggiolini, F.A., Harvey, W.T., et al. (2022). Recurrent inversion polymorphisms in humans associate with genetic instability and genomic disorders. Cell 185, 1986–2005 e1926. 10.1016/j.cell.2022.04.017.

60. Hussin, J.G., Hodgkinson, A., Idaghdour, Y., Grenier, J.C., Goulet, J.P., Gbeha, E., Hip-Ki, E., and Awadalla, P. (2015). Recombination affects accumulation of damaging and disease-associated mutations in human populations. Nat Genet 47, 400–404. 10.1038/ng.3216.

61. Pemberton, T.J., and Szpiech, Z.A. (2018). Relationship between Deleterious Variation, Genomic Autozygosity, and Disease Risk: Insights from The 1000 Genomes Project. Am J Hum Genet 102, 658–675. 10.1016/j.ajhg.2018.02.013.

62. Szpiech, Z.A., Xu, J., Pemberton, T.J., Peng, W., Zollner, S., Rosenberg, N.A., and Li, J.Z. (2013). Long runs of homozygosity are enriched for deleterious variation. Am J Hum Genet 93, 90–102. 10.1016/j.ajhg.2013.05.003.

63. Szpiech, Z.A., Mak, A.C.Y., White, M.J., Hu, D., Eng, C., Burchard, E.G., and Hernandez, R.D. (2019). Ancestry-Dependent Enrichment of Deleterious Homozygotes in Runs of Homozygosity. Am J Hum Genet 105, 747–762. 10.1016/j.ajhg.2019.08.011.

64. Coban-Akdemir, Z., White, J.J., Song, X., Jhangiani, S.N., Fatih, J.M., Gambin, T., Bayram, Y., Chinn, I.K., Karaca, E., Punetha, J., et al. (2018). Identifying Genes Whose Mutant Transcripts Cause Dominant Disease Traits by Potential Gain-of-Function Alleles. Am J Hum Genet 103, 171–187. 10.1016/j.ajhg.2018.06.009.

65. Resnik, P. (1995). Using information content to evaluate semantic similarity in a taxonomy. In: Proceedings of the 14th International Joint Conference on Artificial Intelligence, 448–453.

66. Kohler, S., Doelken, S.C., Mungall, C.J., Bauer, S., Firth, H.V., Bailleul-Forestier, I., Black, G.C., Brown, D.L., Brudno, M., Campbell, J., et al. (2014). The Human Phenotype Ontology project: linking molecular biology and disease through phenotype data. Nucleic Acids Res 42, D966–974. 10.1093/nar/gkt1026.

67. Groza, T., Kohler, S., Moldenhauer, D., Vasilevsky, N., Baynam, G., Zemojtel, T., Schriml, L.M., Kibbe, W.A., Schofield, P.N., Beck, T., et al. (2015). The Human Phenotype Ontology: Semantic Unification of Common and Rare Disease. Am J Hum Genet 97, 111–124. 10.1016/j.ajhg.2015.05.020.

68. Jolly, A., Bayram, Y., Turan, S., Aycan, Z., Tos, T., Abali, Z.Y., Hacihamdioglu, B., Coban Akdemir, Z.H., Hijazi, H., Bas, S., et al. (2019). Exome Sequencing of a Primary Ovarian Insufficiency Cohort Reveals Common Molecular Etiologies for a Spectrum of Disease. J Clin Endocrinol Metab 104, 3049–3067. 10.1210/jc.2019-00248.

69. Zhang, C., Jolly, A., Shayota, B.J., Mazzeu, J.F., Du, H., Dawood, M., Soper, P.C., Ramalho de Lima, A., Ferreira, B.M., Coban-Akdemir, Z., et al. (2022). Novel pathogenic variants and quantitative phenotypic analyses of Robinow syndrome: WNT signaling perturbation and phenotypic variability. HGG Adv 3, 100074. 10.1016/j.xhgg.2021.100074.

70. Herman, I., Jolly, A., Du, H., Dawood, M., Abdel-Salam, G.M.H., Marafi, D., Mitani, T., Calame, D.G., Coban-Akdemir, Z., Fatih, J.M., et al. (2022). Quantitative dissection of multilocus pathogenic variation in an Egyptian infant with severe neurodevelopmental disorder resulting from multiple molecular diagnoses. Am J Med Genet A 188, 735–750. 10.1002/ajmg.a.62565.

71. Lek, M., Karczewski, K.J., Minikel, E.V., Samocha, K.E., Banks, E., Fennell, T., O’Donnell-Luria, A.H., Ware, J.S., Hill, A.J., Cummings, B.B., et al. (2016). Analysis of protein-coding genetic variation in 60,706 humans. Nature 536, 285–291. 10.1038/nature19057.

72. Gambin, T., Jhangiani, S.N., Below, J.E., Campbell, I.M., Wiszniewski, W., Muzny, D.M., Staples, J., Morrison, A.C., Bainbridge, M.N., Penney, S., et al. (2015). Secondary findings and carrier test frequencies in a large multiethnic sample. Genome Med 7, 54. 10.1186/s13073-015-0171-1.

73. Bayram, Y., Karaca, E., Coban Akdemir, Z., Yilmaz, E.O., Tayfun, G.A., Aydin, H., Torun, D., Bozdogan, S.T., Gezdirici, A., Isikay, S., et al. (2016). Molecular etiology of arthrogryposis in multiple families of mostly Turkish origin. J Clin Invest 126, 762–778. 10.1172/JCI84457.

74. Narasimhan, V.M., Rahbari, R., Scally, A., Wuster, A., Mason, D., Xue, Y., Wright, J., Trembath, R.C., Maher, E.R., van Heel, D.A., et al. (2017). Estimating the human mutation rate from autozygous segments reveals population differences in human mutational processes. Nat Commun 8, 303. 10.1038/s41467-017-00323-y.

75. Zollo, M., Ahmed, M., Ferrucci, V., Salpietro, V., Asadzadeh, F., Carotenuto, M., Maroofian, R., Al-Amri, A., Singh, R., Scognamiglio, I., et al. (2017). *PRUNE* is crucial for normal brain development and mutated in microcephaly with neurodevelopmental impairment. Brain 140, 940–952. 10.1093/brain/awx014.

76. Nistala, H., Dronzek, J., Gonzaga-Jauregui, C., Chim, S.M., Rajamani, S., Nuwayhid, S., Delgado, D., Burke, E., Karaca, E., Franklin, M.C., et al. (2021). NMIHBA results from hypomorphic *PRUNE1* variants that lack short-chain exopolyphosphatase activity. Hum Mol Genet 29, 3516–3531. 10.1093/hmg/ddaa237.

77. Karaca, E., Weitzer, S., Pehlivan, D., Shiraishi, H., Gogakos, T., Hanada, T., Jhangiani, S.N., Wiszniewski, W., Withers, M., Campbell, I.M., et al. (2014). Human *CLP1* mutations alter tRNA biogenesis, affecting both peripheral and central nervous system function. Cell 157, 636–650. 10.1016/j.cell.2014.02.058.

78. Schaffer, A.E., Eggens, V.R., Caglayan, A.O., Reuter, M.S., Scott, E., Coufal, N.G., Silhavy, J.L., Xue, Y., Kayserili, H., Yasuno, K., et al. (2014). *CLP1* founder mutation links tRNA splicing and maturation to cerebellar development and neurodegeneration. Cell 157, 651–663. 10.1016/j.cell.2014.03.049.

79. Yuan, B., Pehlivan, D., Karaca, E., Patel, N., Charng, W.L., Gambin, T., Gonzaga-Jauregui, C., Sutton, V.R., Yesil, G., Bozdogan, S.T., et al. (2015). Global transcriptional disturbances underlie Cornelia de Lange syndrome and related phenotypes. J Clin Invest 125, 636–651. 10.1172/JCI77435.

80. Rainger, J., Pehlivan, D., Johansson, S., Bengani, H., Sanchez-Pulido, L., Williamson, K.A., Ture, M., Barker, H., Rosendahl, K., Spranger, J., et al. (2014). Monoallelic and biallelic mutations in *MAB21L2* cause a spectrum of major eye malformations. Am J Hum Genet 94, 915–923. 10.1016/j.ajhg.2014.05.005.

81. Monies, D., Abouelhoda, M., Assoum, M., Moghrabi, N., Rafiullah, R., Almontashiri, N., Alowain, M., Alzaidan, H., Alsayed, M., Subhani, S., et al. (2019). Lessons Learned from Large-Scale, First-Tier Clinical Exome Sequencing in a Highly Consanguineous Population. Am J Hum Genet 104, 1182–1201. 10.1016/j.ajhg.2019.04.011.

82. Carvalho, C.M.B., Coban-Akdemir, Z., Hijazi, H., Yuan, B., Pendleton, M., Harrington, E., Beaulaurier, J., Juul, S., Turner, D.J., Kanchi, R.S., et al. (2019). Interchromosomal template-switching as a novel molecular mechanism for imprinting perturbations associated with Temple syndrome. Genome Med 11, 25. 10.1186/s13073-019-0633-y.

83. Lupski, J.R., Liu, P., Stankiewicz, P., Carvalho, C.M.B., and Posey, J.E. (2020). Clinical genomics and contextualizing genome variation in the diagnostic laboratory. Expert Rev Mol Diagn 20, 995–1002. 10.1080/14737159.2020.1826312.

84. Castel, S.E., Cervera, A., Mohammadi, P., Aguet, F., Reverter, F., Wolman, A., Guigo, R., Iossifov, I., Vasileva, A., and Lappalainen, T. (2018). Modified penetrance of coding variants by cis-regulatory variation contributes to disease risk. Nat Genet 50, 1327–1334. 10.1038/s41588-018-0192-y.

85. Yang, N., Wu, N., Zhang, L., Zhao, Y., Liu, J., Liang, X., Ren, X., Li, W., Chen, W., Dong, S., et al. (2019). *TBX6* compound inheritance leads to congenital vertebral malformations in humans and mice. Hum Mol Genet 28, 539–547. 10.1093/hmg/ddy358.

86. Liu, J., Wu, N., Deciphering Disorders Involving, S., study, C.O., Yang, N., Takeda, K., Chen, W., Li, W., Du, R., Liu, S., et al. (2019). *TBX6*-associated congenital scoliosis (TACS) as a clinically distinguishable subtype of congenital scoliosis: further evidence supporting the compound inheritance and *TBX6* gene dosage model. Genet Med 21, 1548–1558. 10.1038/s41436-018-0377-x.

87. Ren, X., Yang, N., Wu, N., Xu, X., Chen, W., Zhang, L., Li, Y., Du, R.Q., Dong, S., Zhao, S., et al. (2020). Increased *TBX6* gene dosages induce congenital cervical vertebral malformations in humans and mice. J Med Genet 57, 371–379. 10.1136/jmedgenet-2019-106333.

88. Duan, R., Hijazi, H., Gulec, E.Y., Eker, H.K., Costa, S.R., Sahin, Y., Ocak, Z., Isikay, S., Ozalp, O., Bozdogan, S., et al. (2022). Developmental genomics of limb malformations: Allelic series in association with gene dosage effects contribute to the clinical variability. HGG Adv 3, 100132. 10.1016/j.xhgg.2022.100132.

89. Yuan, B., Schulze, K.V., Assia Batzir, N., Sinson, J., Dai, H., Zhu, W., Bocanegra, F., Fong, C.T., Holder, J., Nguyen, J., et al. (2022). Sequencing individual genomes with recurrent genomic disorder deletions: an approach to characterize genes for autosomal recessive rare disease traits. Genome Med 14, 113. 10.1186/s13073-022-01113-y.

90. Gambin, T., Akdemir, Z.C., Yuan, B., Gu, S., Chiang, T., Carvalho, C.M.B., Shaw, C., Jhangiani, S., Boone, P.M., Eldomery, M.K., et al. (2017). Homozygous and hemizygous CNV detection from exome sequencing data in a Mendelian disease cohort. Nucleic Acids Res 45, 1633–1648. 10.1093/nar/gkw1237.

91. Yuan, B., Wang, L., Liu, P., Shaw, C., Dai, H., Cooper, L., Zhu, W., Anderson, S.A., Meng, L., Wang, X., et al. (2020). CNVs cause autosomal recessive genetic diseases with or without involvement of SNV/indels. Genet Med 22, 1633–1641. 10.1038/s41436-020-0864-8.

92. Dharmadhikari, A.V., Ghosh, R., Yuan, B., Liu, P., Dai, H., Al Masri, S., Scull, J., Posey, J.E., Jiang, A.H., He, W., et al. (2019). Copy number variant and runs of homozygosity detection by microarrays enabled more precise molecular diagnoses in 11,020 clinical exome cases. Genome Med 11, 30. 10.1186/s13073-019-0639-5.

93. Karolak, J.A., Vincent, M., Deutsch, G., Gambin, T., Cogne, B., Pichon, O., Vetrini, F., Mefford, H.C., Dines, J.N., Golden-Grant, K., et al. (2019). Complex Compound Inheritance of Lethal Lung Developmental Disorders Due to Disruption of the TBX-FGF Pathway. Am J Hum Genet 104, 213–228. 10.1016/j.ajhg.2018.12.010.

94. Lupski, J.R., Belmont, J.W., Boerwinkle, E., and Gibbs, R.A. (2011). Clan genomics and the complex architecture of human disease. Cell 147, 32–43. 10.1016/j.cell.2011.09.008.

95. Lupski, J.R. (2021). Clan genomics: From OMIM phenotypic traits to genes and biology. Am J Med Genet A 185, 3294–3313. 10.1002/ajmg.a.62434.

96. Lupski, J.R. (2022). Biology in balance: human diploid genome integrity, gene dosage, and genomic medicine. Trends Genet 38, 554–571. 10.1016/j.tig.2022.03.001.

