## Supplementary Figures for "The impact of the Turkish (TK) population variome on the genomic architecture of rare disease traits"

**Supplementary Figure 1**

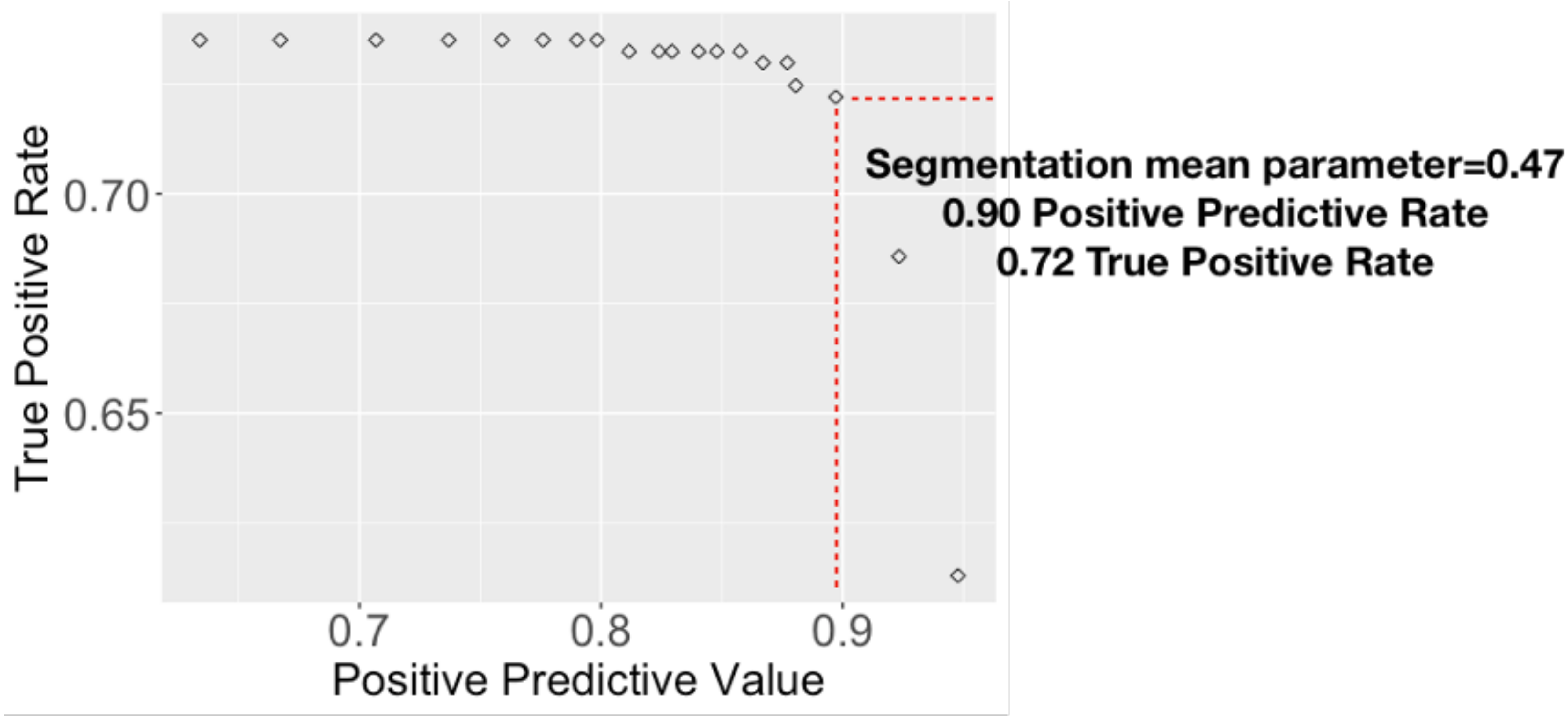

**Supplementary Figure 2**

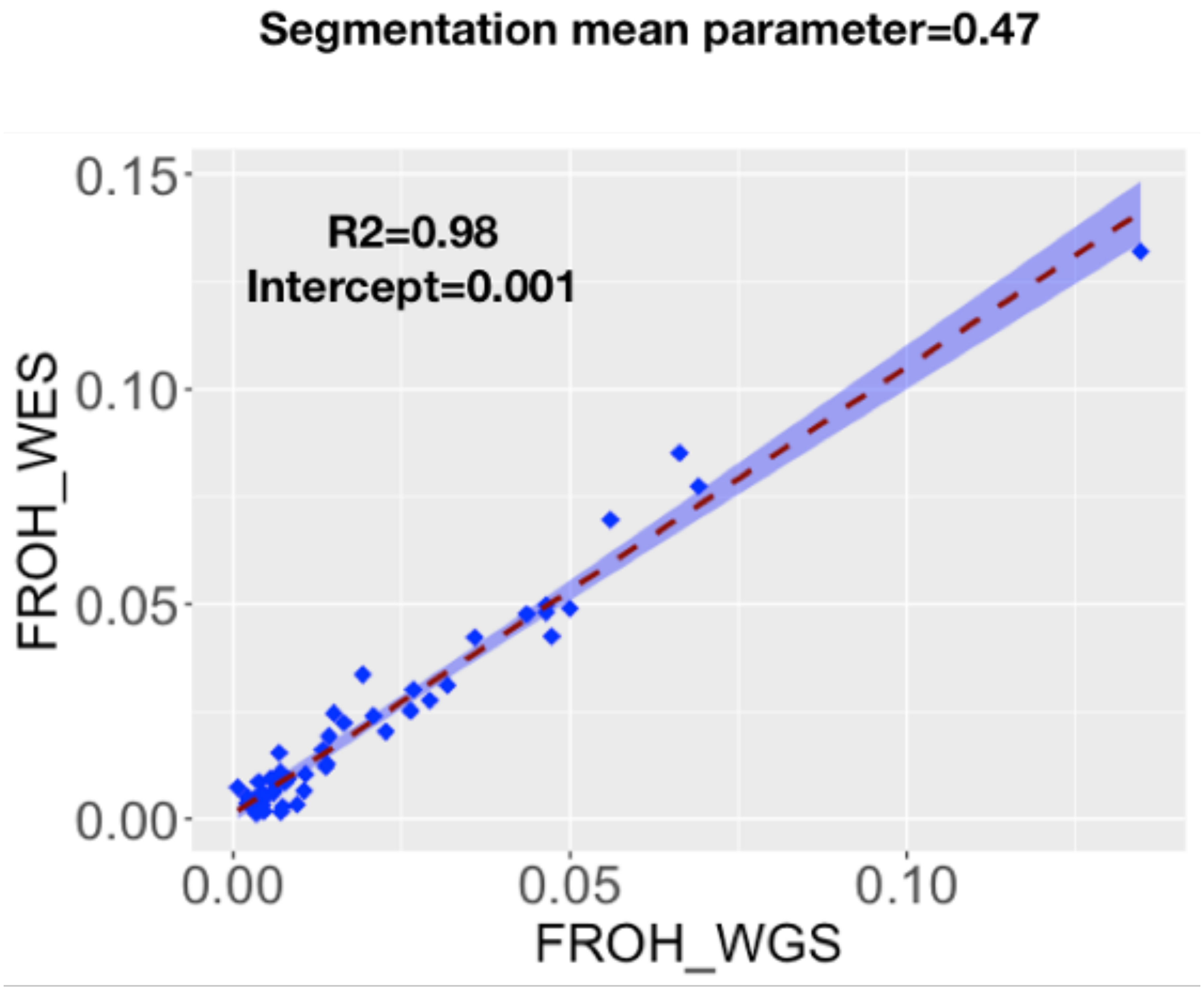

**Supplementary Figure 3**

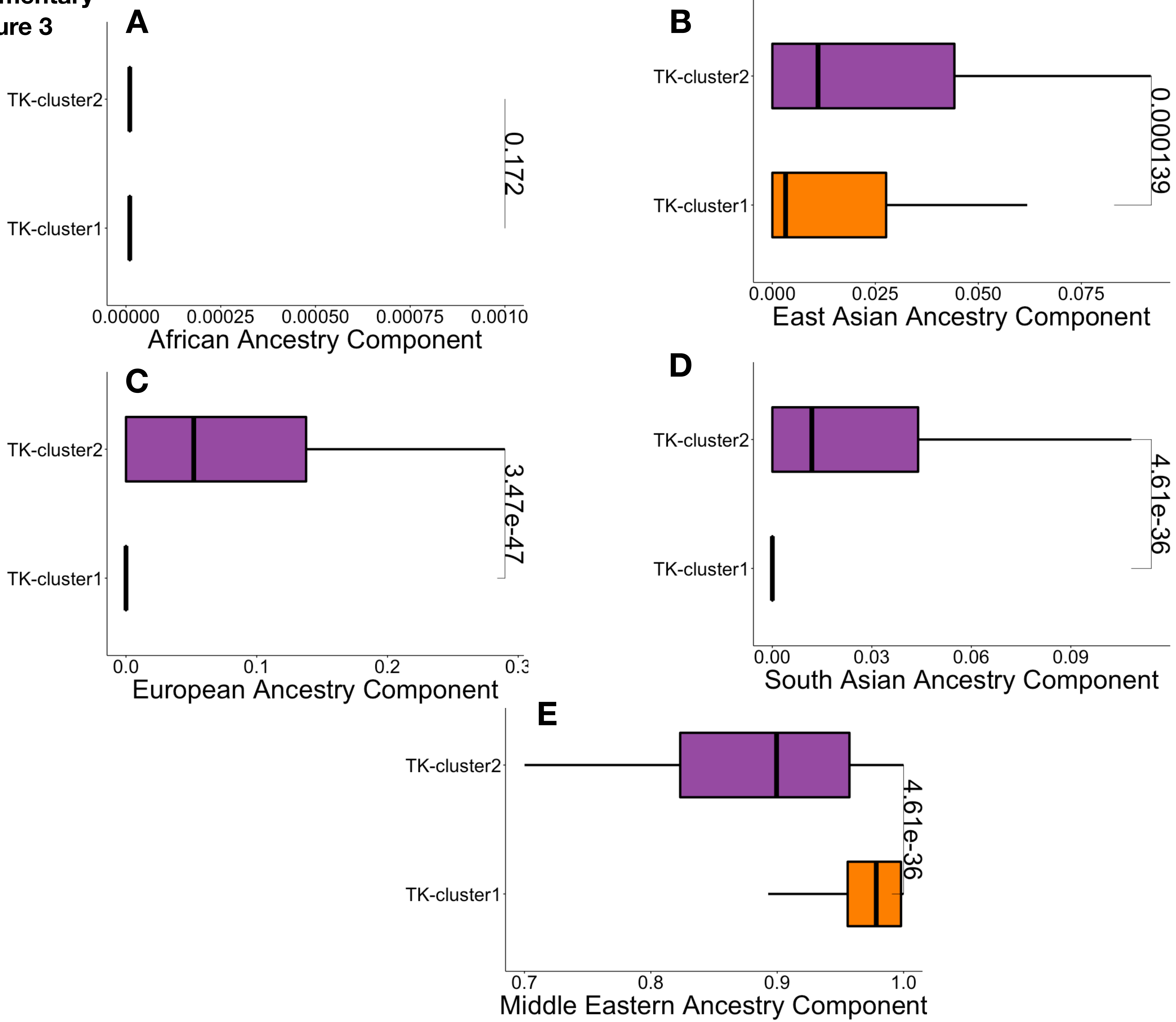

Supplementary Figure 4

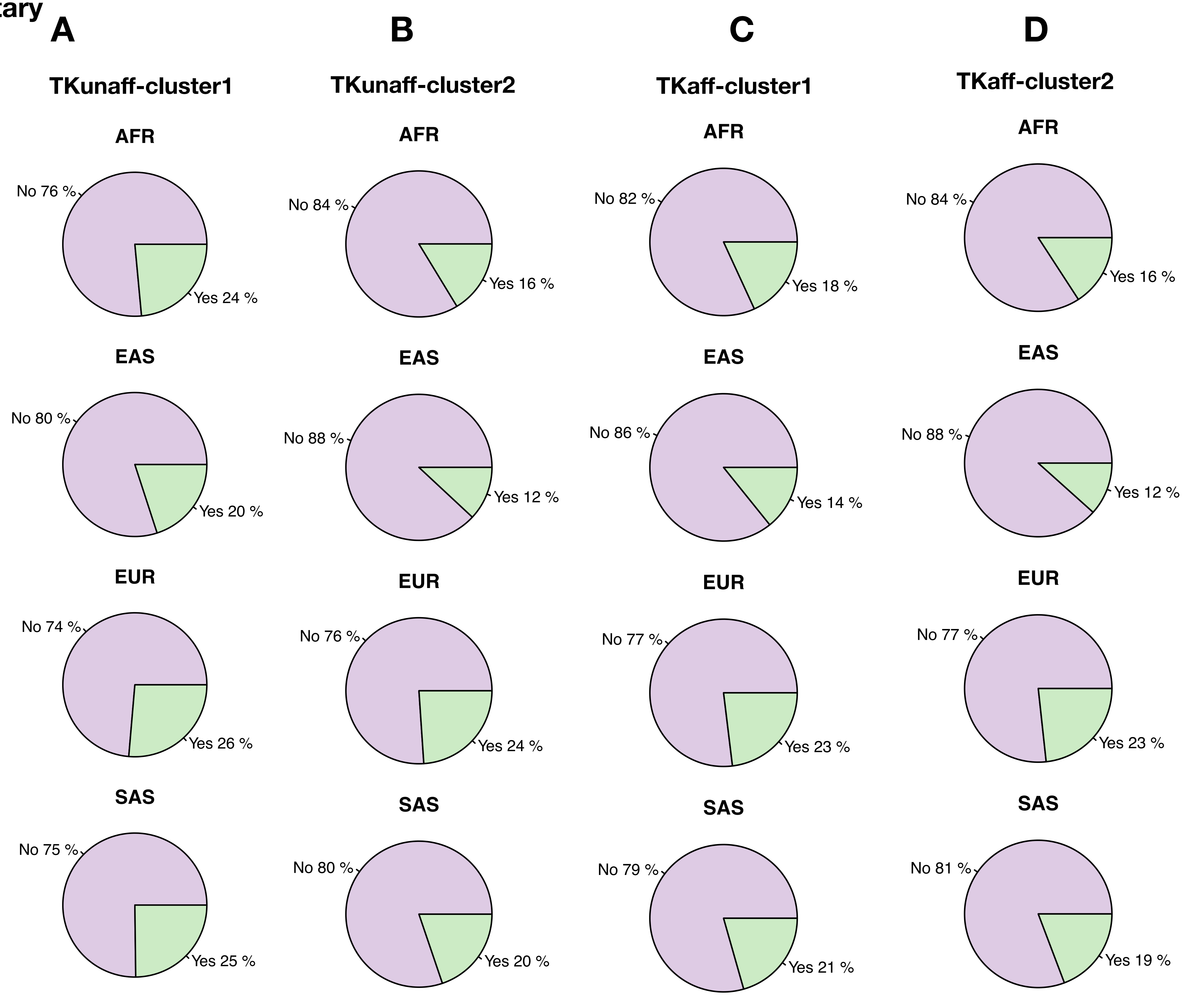

**Supplementary Figure 5**

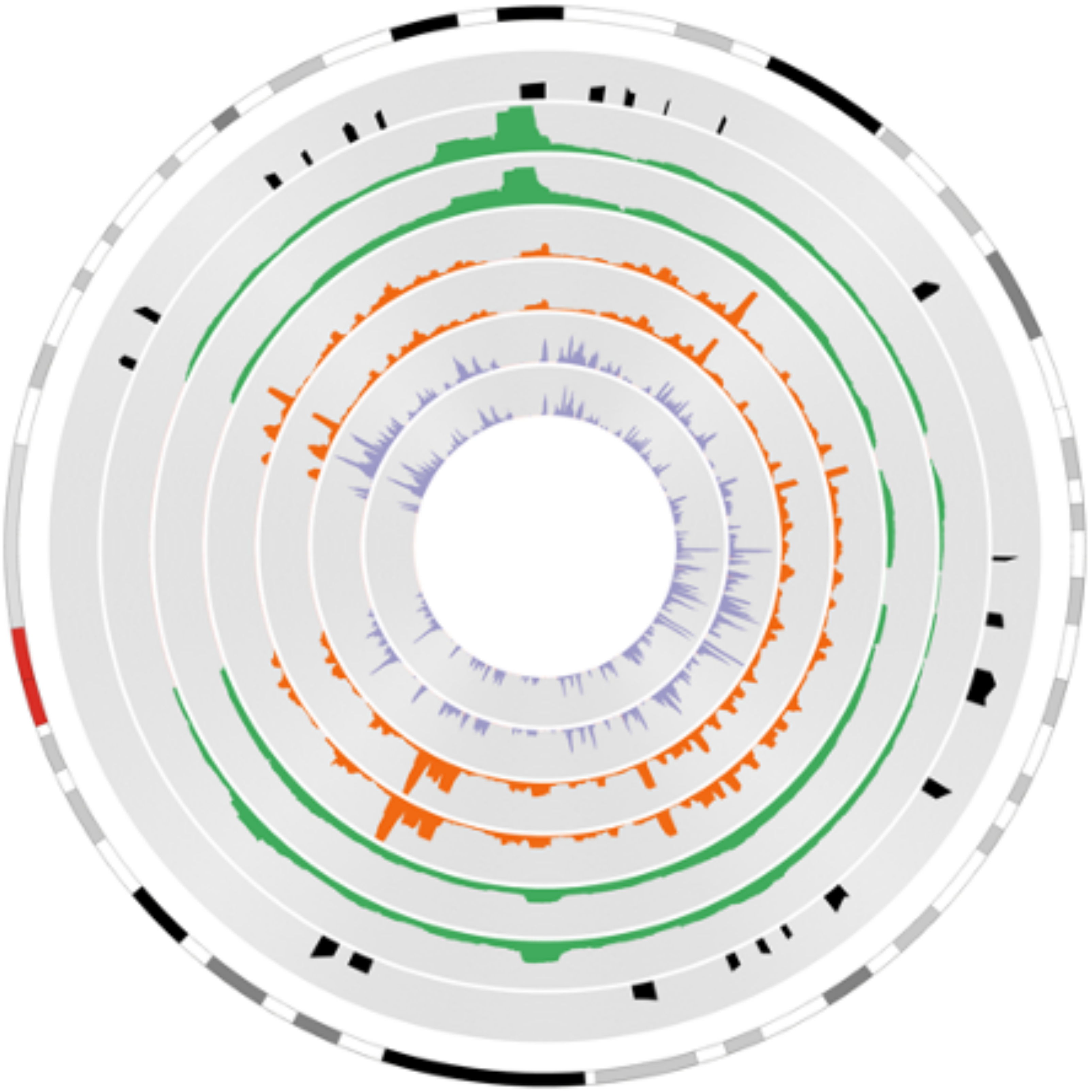

### Supplemental Figure 6

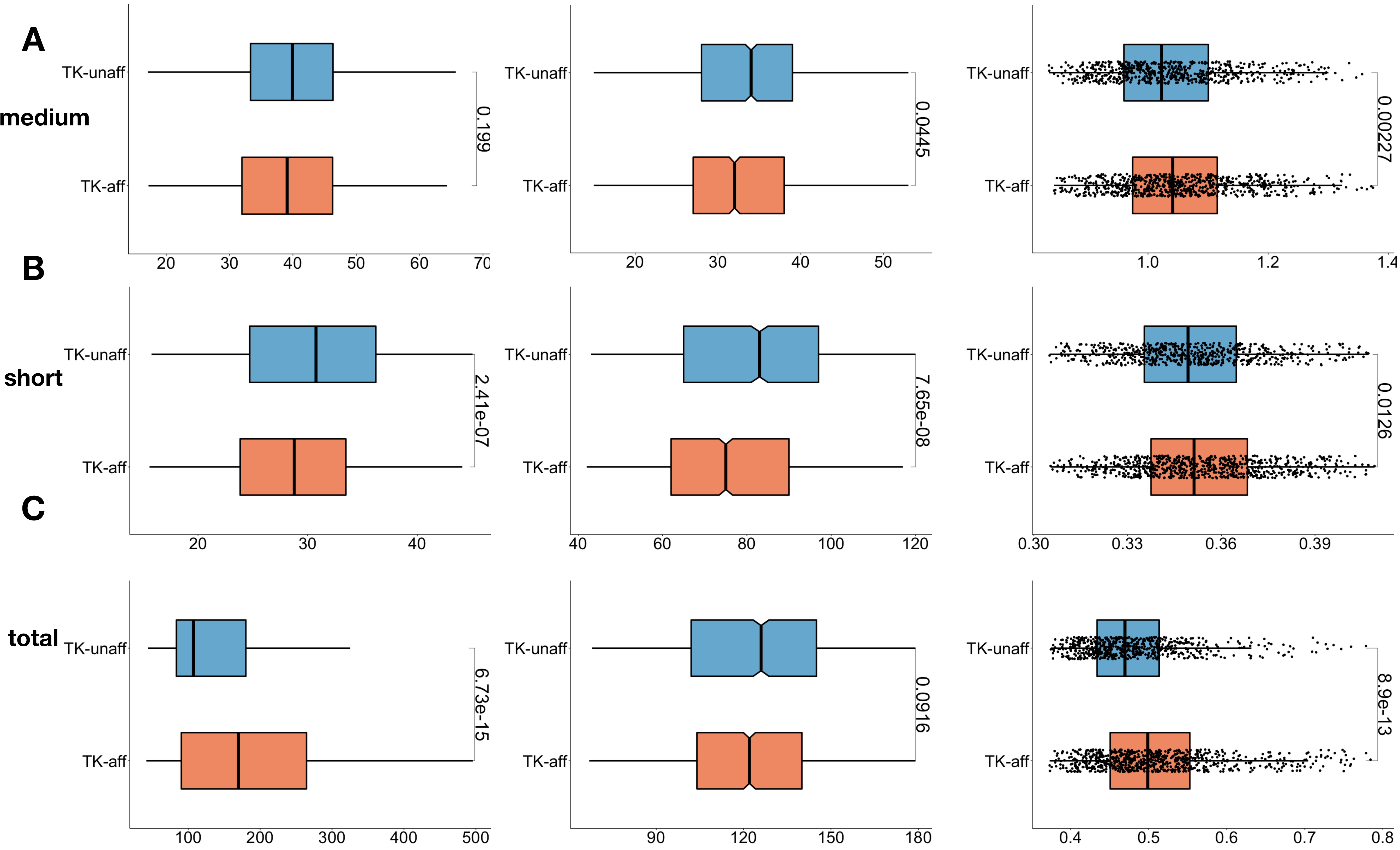

**Supplementary Figure 7**

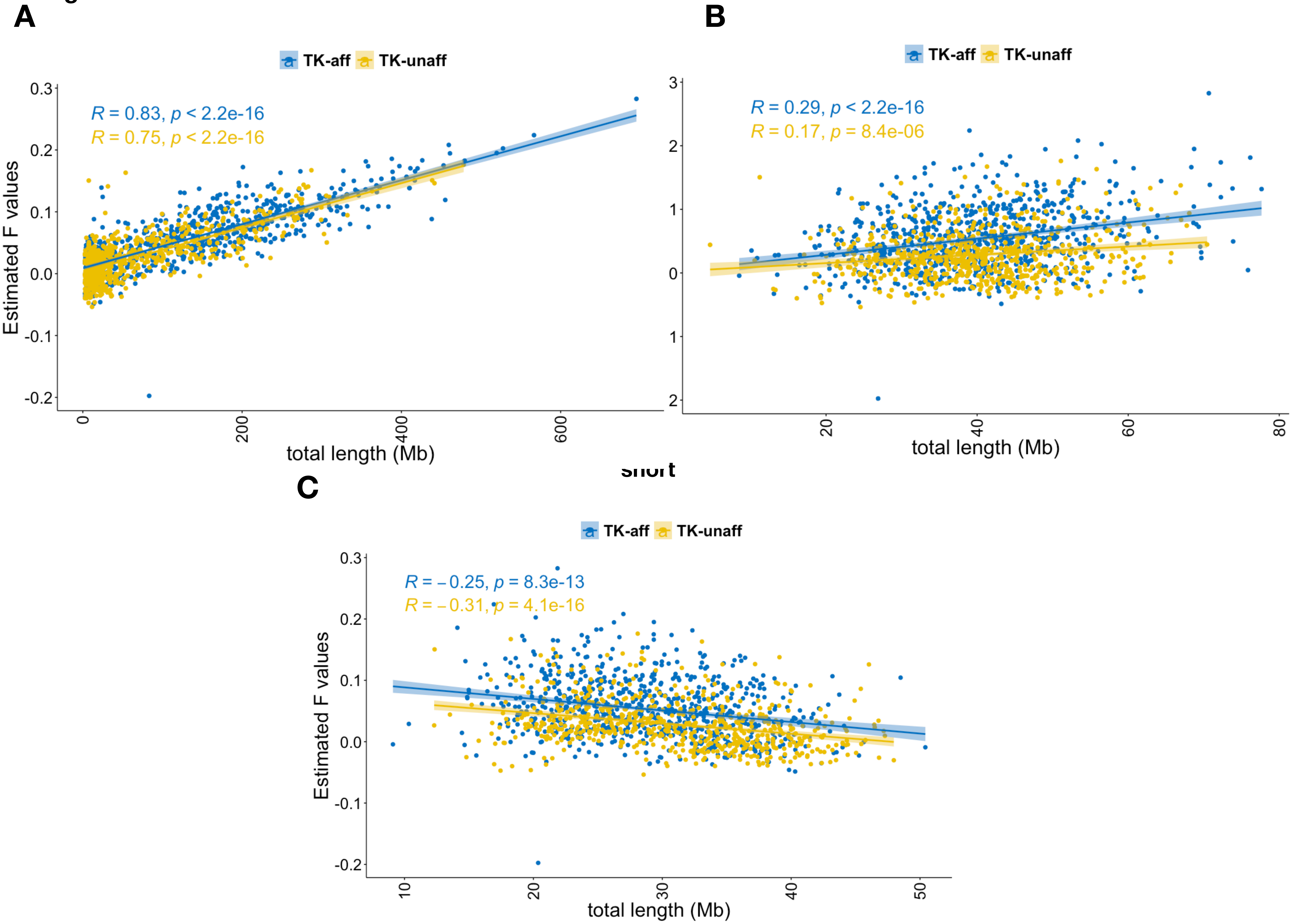

### Supplementary Figure 8

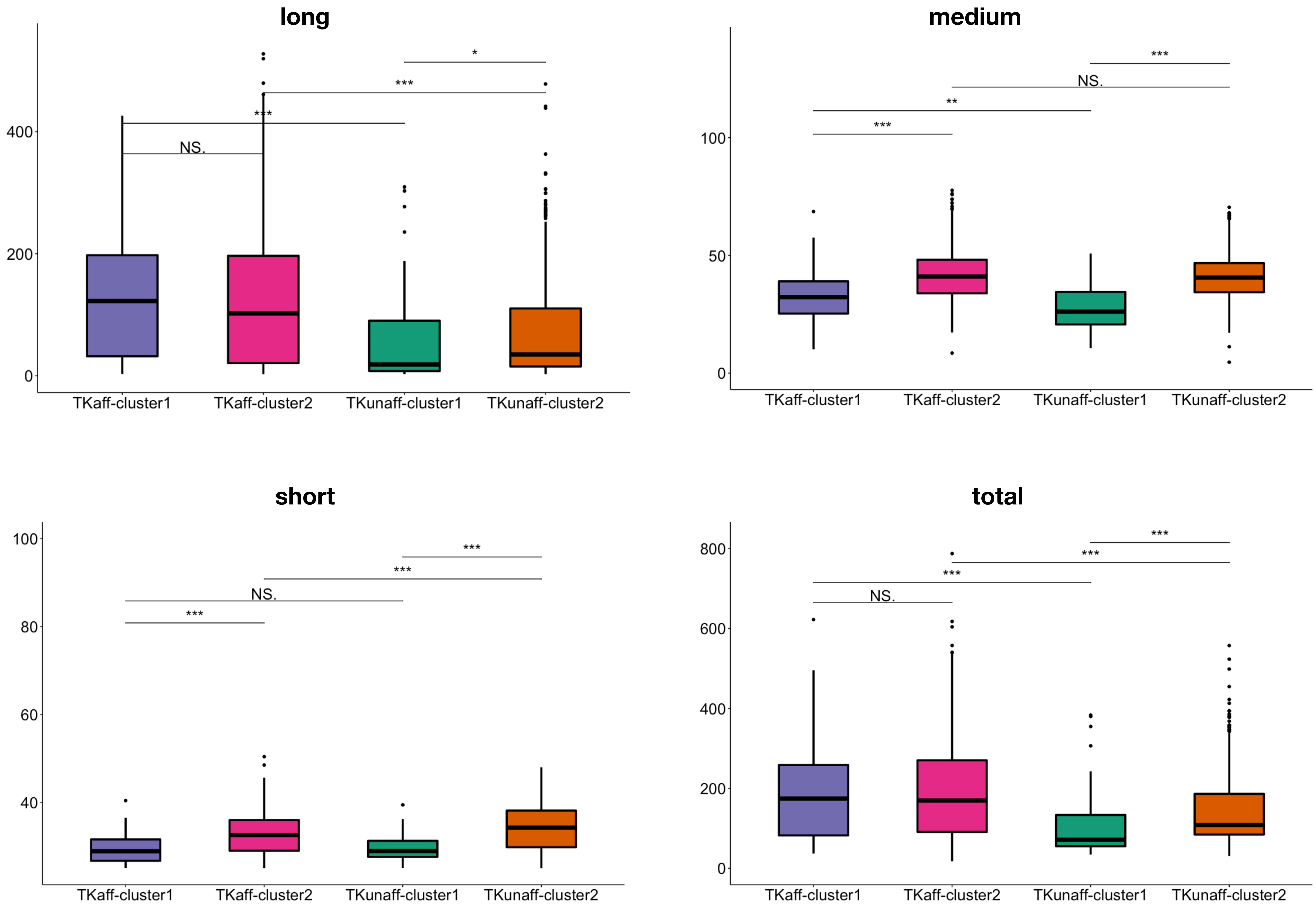

Supplementary Figure 9

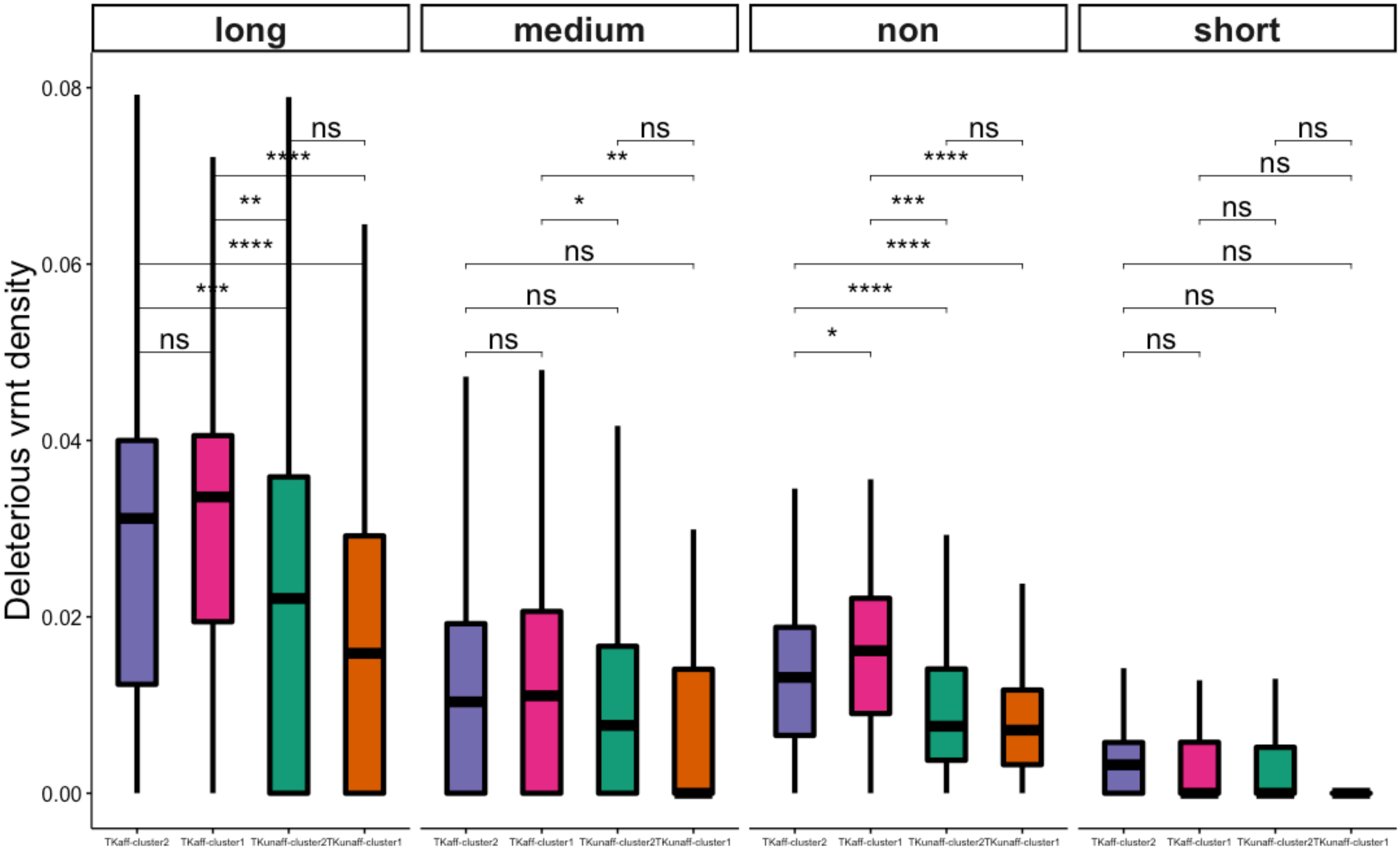

#### Supplementary Figure 10

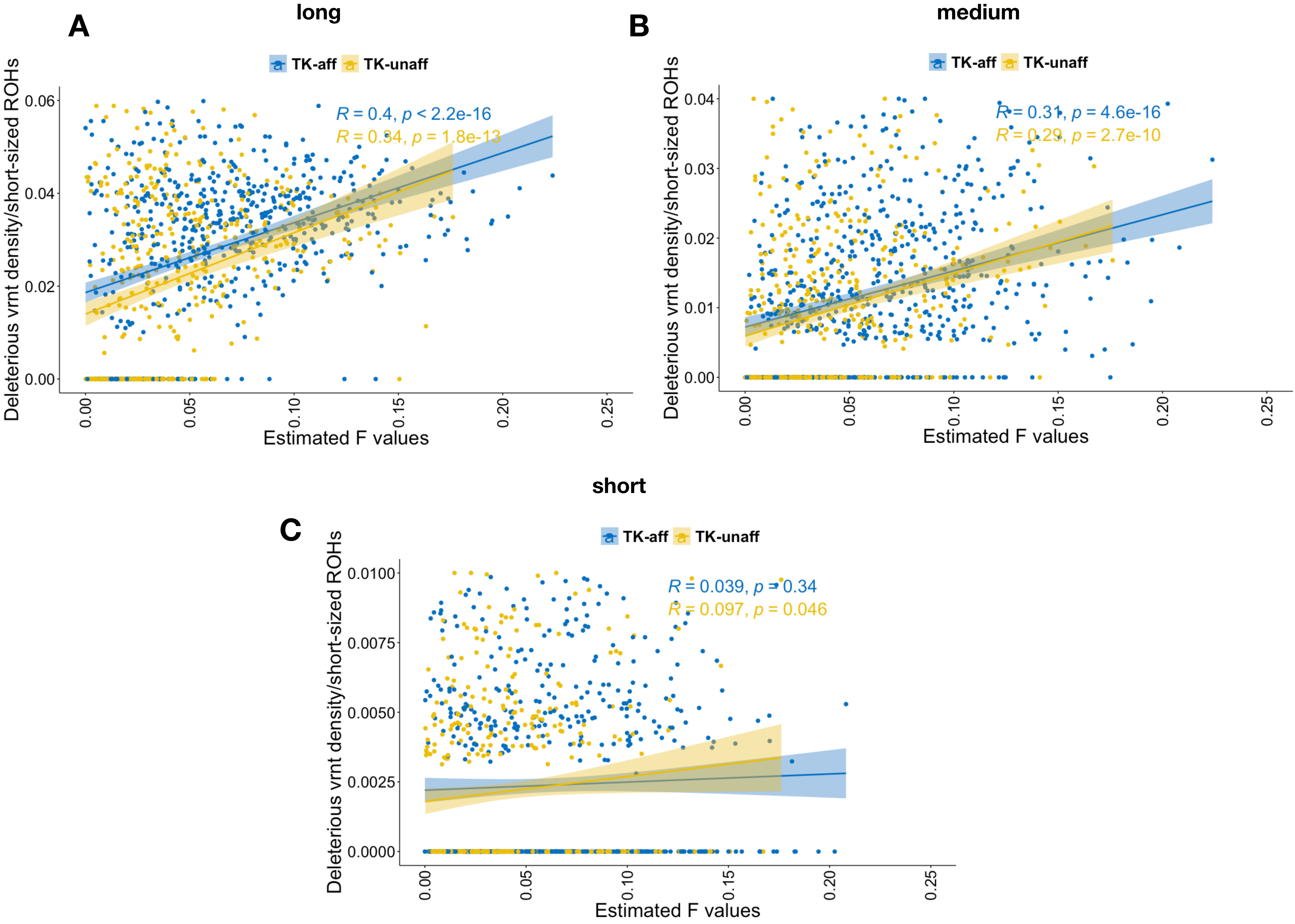

**Supplementary Figure 11**

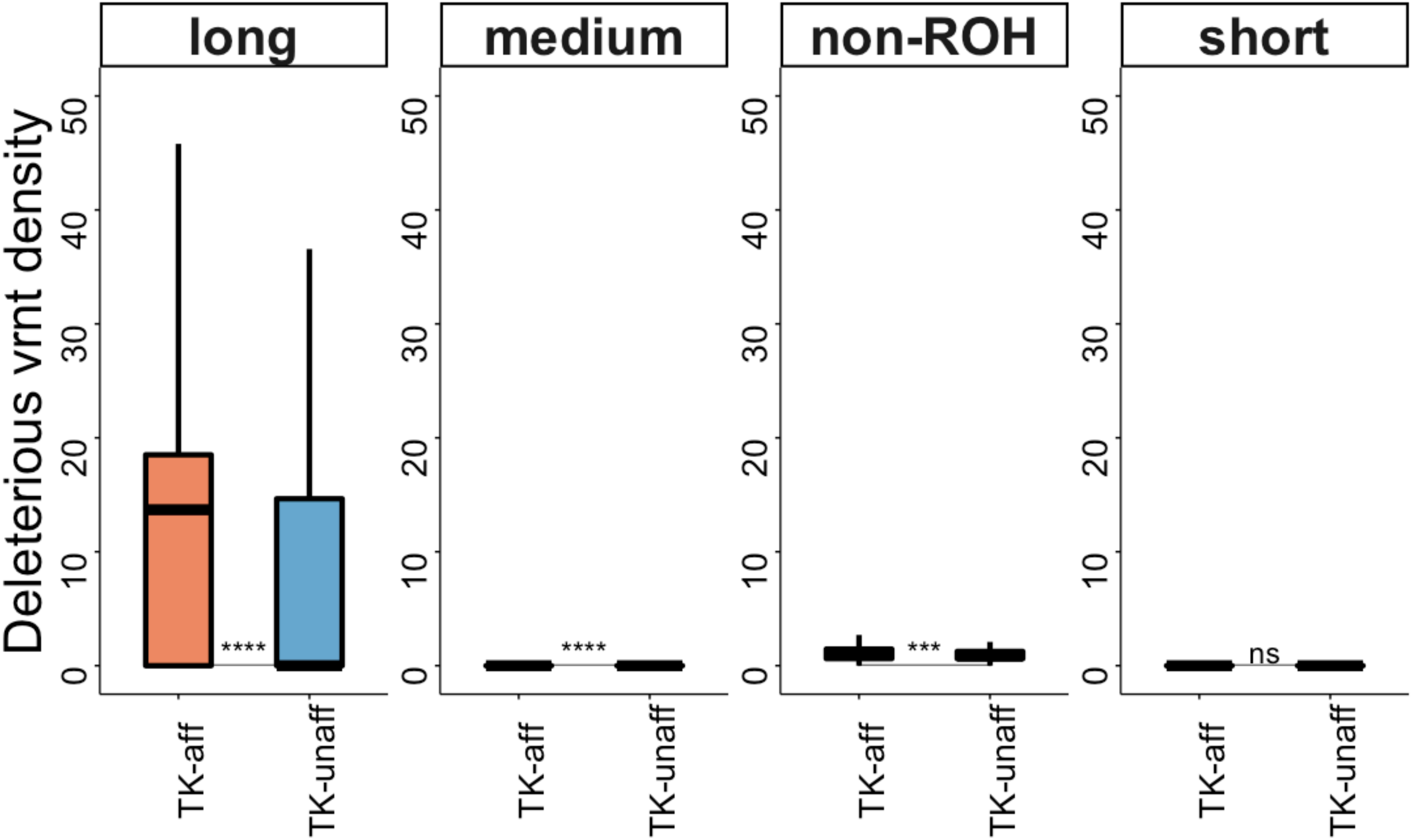
